# Structural and functional studies of the RBPJ-SHARP complex reveal conserved corepressor binding site

**DOI:** 10.1101/273615

**Authors:** Zhenyu Yuan, Bradley D. VanderWielen, Benedetto Daniele Giaimo, Leiling Pan, Courtney E. Collins, Aleksandra Turkiewicz, Kerstin Hein, Franz Oswald, Tilman Borggrefe, Rhett A. Kovall

**Affiliations:** Department of Molecular Genetics, Biochemistry and Microbiology, University of Cincinnati College of Medicine, Cincinnati, Ohio, USA; Institute of Biochemistry, University of Giessen, Giessen, Germany; Department of Internal Medicine I, Center for Internal Medicine, University Medical Center Ulm, 89081 Ulm, Germany.

## Abstract

The Notch pathway is a conserved signaling mechanism that is essential for cell fate decisions during pre and postnatal development. Dysregulated signaling underlies the pathophysiology of numerous human diseases, most notably T-cell acute lymphoblastic leukemia. Receptor-ligand interactions result in changes in gene expression, which are regulated by the transcription factor CSL. CSL forms a complex with the intracellular domain of the Notch receptor and the transcriptional coactivator Mastermind, which is required to activate transcription of all Notch target genes. CSL can also function as repressor by interacting with corepressor proteins, *e.g*. SHARP in mammals and Hairless in *Drosophila melanogaster;* however, its role as a transcriptional repressor is not well understood. Here we determine the high-resolution structure of RBPJ, the mouse CSL ortholog, bound to the corepressor SHARP and DNA, which reveals a new mode of corepressor binding to CSL and an interesting example for how ligand binding sites evolve in proteins. Based on the structure, we designed and tested a number of mutants in biophysical, biochemical, and cellular assays to characterize the role of RBPJ as a repressor of Notch target genes. Our cellular studies clearly demonstrate that RBPJ mutants that are deficient for binding SHARP are incapable of repressing transcription from genes responsive to Notch signaling. Altogether, our structure-function studies of the RBPJ-SHARP corepressor complex bound to DNA provide significant insights into the repressor function of RBPJ and identify a new binding pocket on RBPJ that could be targeted for therapeutic benefit.

## Introduction

The Notch pathway is a highly conserved cell-to-cell signaling mechanism that is indispensable for cell fate decisions during prenatal development and postnatal tissue homeostasis.^1,2^ Dysregulated signaling underlies the pathogenesis of many human diseases, including certain types of cancer, congenital defects, and cardiovascular disease^3^. Given its association with human disease, there have been extensive efforts to identify reagents that pharmaceutically modulate the Notch pathway with most efforts focused on modalities that curtail overactive Notch signaling^4,5^. However, there is a need to identify components in the Notch pathway that when pharmacologically targeted result in upregulated signaling to treat diseases associated with insufficient Notch signaling^3^, such as myeloid leukemia, head and neck squamous cell carcinoma, bicuspid aortic valve disease, and Alagille syndrome.

Notch signaling is initiated when Notch receptors on one cell interact with a DSL (Delta, Serrate, Lag-2) ligand on a neighboring cell (Figure 1A)^6^. Extracellular Notch-DSL interactions result in proteolytic cleavage of the receptor, which frees the intracellular domain of Notch (NICD) from the cell membrane, allowing NICD to migrate into the nucleus. Nuclear NICD directly binds the transcription factor CSL [CBF1/RBPJ, Su(H), Lag-1] and the CSL-NICD complex recruits a member of the Mastermind (MAM) family of transcriptional coactivators [Mastermind-like (MAML1-3) in mammals], resulting in the transcriptional activation of Notch target genes (Figure 1A)^7^. CSL can also function as a repressor by interacting with corepressor proteins, such as SHARP [SMRT/HDAC1-associated repressor protein, *also known as* MINT (Msx2-interacting nuclear target) or SPEN (split ends)]^8,9^, Hairless in *Drosophila melanogaster^10^*, FHL1 (Four and a half LIM domains 1, *also known as* KyoT2)^11^, L3MBTL3 (lethal 3 malignant brain tumor-like 3)^12^, and RITA1 (RBPJ-interacting and tubulin associated)^13,14^. Corepressors are part of large higher order transcriptional repression complexes that contain histone-modifying activity, *e.g*. histone deacetylase or histone demethylase, which convert chromatin into a transcriptionally repressed state (Figure 1A)^7^.

**Figure 1.**
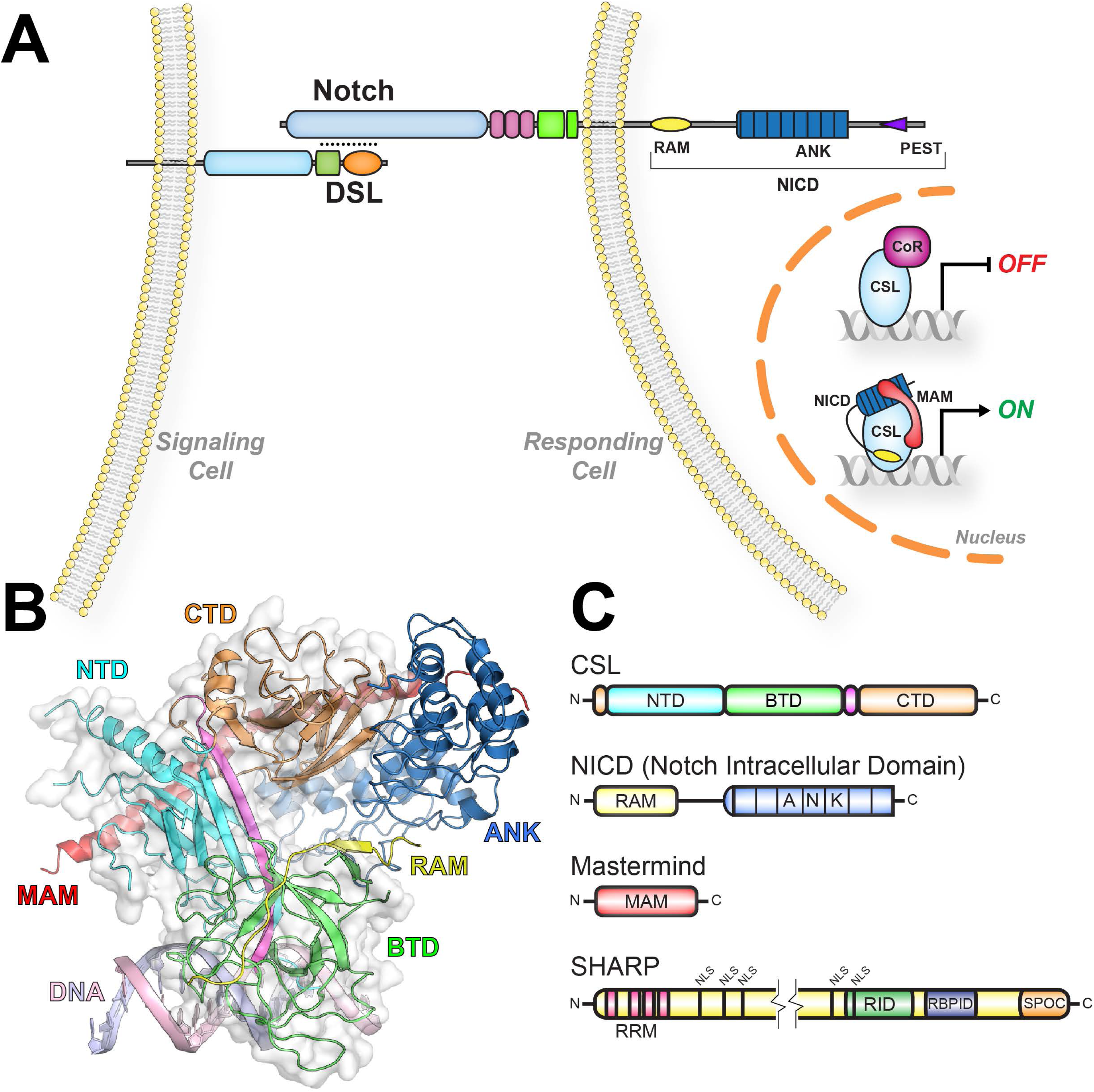
<u>Overview of Notch signaling</u>. (A) Notch signaling occurs between neighboring cells, in which extracellular Notch-DSL interactions initiate signaling, resulting in the cleavage of Notch and release of its intracellular domain (NICD) from the cell membrane that subsequently transits to the nucleus. In the absence of Notch signaling, CSL can bind corepressors (CoR) to repress transcription from Notch target genes. Upon pathway activation, NICD and Mastermind (MAM) form a ternary complex with CSL that activates transcription from Notch target genes. (B) Structure of the CSL-NICD-MAM ternary complex bound to DNA (PDB ID: 2FO1). The structural core of CSL is composed of three domains: NTD (N-terminal domain), BTD (β-trefoil domain), and CTD (C-terminal domain), which are colored cyan, green, and orange, respectively. A β-strand that makes hydrogen-bonding interactions with all three domains is colored magenta. The RAM and ANK domains of NICD are colored yellow and blue, respectively. MAM and DNA are colored red and light pink/blue, respectively. (C) Domain schematics of CSL, NICD, and MAM are colored similarly to the structure. SHARP is a multidomain transcriptional coregulator that contains N-terminal RRM (RNA Recognition Motif) domains, multiple NLS (Nuclear Localization Sequence), a RID (Receptor Interaction Domain), a RBPID (RBPJ Interacting Domain), and a C-terminal SPOC (Spen Paralog and Ortholog C-terminal) domain.

Crystal structures of Notch transcription complexes have revealed that all CSL proteins contain a conserved structural core composed of three domains termed NTD (N-terminal domain), BTD (β-trefoil domain), and CTD (C-terminal domain) (Figure 1B,C)^15–17^. The NTD and CTD are immunoglobulin (Ig) domains that are structurally similar to the Rel homology region (RHR) of transcription factors, such as NF-κB and NFAT (Nuclear factor of activated T-cells), whereas the fold of the BTD is related to cytokine and growth factor structures, such as Interleukin1 and FGF (Fibroblast Growth Factor)^17^. The NTD and BTD form an electropositive surface that recognizes nucleotides in the major and minor grooves of DNA, respectively^17^. In the transcriptionally active CSL-NICD-MAM ternary complex bound to DNA structure (Figure 1B,C), the RAM and ANK domains of NICD bind the BTD and CTD domains of CSL, respectively, and MAM forms an elongated bent helical structure that binds the CTD-ANK interface and the NTD of CSL^15,16^. In addition to the CSL-NICD-MAM activator complex, several CSL-corepressor structures have been determined, including the *Drosophila* corepressor Hairless bound to Su(H) (the fly CSL ortholog)^18^, as well as FHL1 and RITA1 bound to RBPJ (mouse/human CSL ortholog)^13,19^. These studies reveal that the binding of Hairless to the CTD of Su(H) produces a large conformational change in CTD^18^; whereas, FHL1 and RITA1 have regions similar to the RAM domain of NICD and bind the BTD of RBPJ in a structurally similar manner^13,19^.

SHARP is a large multidomain transcriptional coregulator protein that has folded functional domains separated by long regions that are intrinsically disordered. Notably, SHARP has no sequence similarity with any of the previously identified corepressors that bind CSL (Figure 1C). SHARP was originally identified in yeast two-hybrid screens for factors that interact with SMRT/NCoR and the transcription factor MSX2^20,21^. SHARP has traditionally been thought of as a corepressor, because it binds SMRT/NCoR through its C-terminal SPOC domain and represses transcription^9,20,22,23^; however, it has also been shown that through its SPOC domain SHARP can also recruit the KMT2D coactivator complex to Notch target genes^24^. More recently, it has been demonstrated that the N-terminal RRM (RNA recognition motif) domains of SHARP bind the lncRNA Xist and this interaction is important for X chromosome inactivation^25,26^. Previously, we defined a region in SHARP that binds RBPJ, termed RBPID (RBPJ-interacting domain, Figure 1C) and showed that RBPJ and SHARP form a high affinity complex^9,27^.

Here we determine the X-ray structure of the RBPJ-SHARP corepressor complex bound to DNA. We identify structure-based mutants that are essential for RBPJ-mediated repression and characterize these mutants both *in vitro* and in cellular assays. Taken together, our studies reveal the conserved interface of the RBPJ/SHARP corepressor complex, which provides molecular insights into RBPJ repressor function and identify a potential site on RBPJ that could be pharmacologically targeted to upregulate Notch signaling.

## Results

### Structure determination of the RBPJ/SHARP/DNA complex

In order to determine the X-ray structure of the RBPJ/SHARP corepressor complex bound to DNA, we purified recombinant RBPJ and SHARP proteins from bacteria, corresponding to the structural core of RBPJ (residues 53–474) and the RBPJ-interacting domain (RBPID) of SHARP (residues 2776–2820), formed complexes with a 13mer oligomeric DNA duplex with single-stranded overhangs, containing a single RBPJ binding site, and screened the RBPJ/SHARP/DNA complex for crystallization conditions. While we were able to isolate crystallization conditions for the complex, despite extensive optimization efforts, we were unable to produce diffraction quality crystals amenable for structural analysis. Therefore, we produced an MBP (maltose binding protein) fusion protein with the RBPID of SHARP (MBP-SHARP), *i.e*. fixed-arm carrier approach^28^, in which the MBP moiety also has surface entropy reduction mutations engineered into it, in order to identify new crystallization conditions for the RBPJ/MBP-SHARP/DNA complex. This strategy successfully led to crystals that nominally diffract to 2.8 Å resolution and belong to the space group P2_1_ (a=54.5 Å, b=231.6 Å, c=90.3 Å, and β=99.88°) (Table 1). We demonstrated that our MBP-SHARP construct binds RBPJ similarly as the native SHARP construct, albeit with somewhat weaker affinity (Figure S1B). This is likely due to the close crystal contacts between MBP and RBPJ required for crystallization of the complex. The RBPJ/MBP-SHARP/DNA complex structure was solved by molecular replacement and refined to a final R factor and free R factor of 19.4% and 22.8%, respectively (Table 1).

**Table 1:**
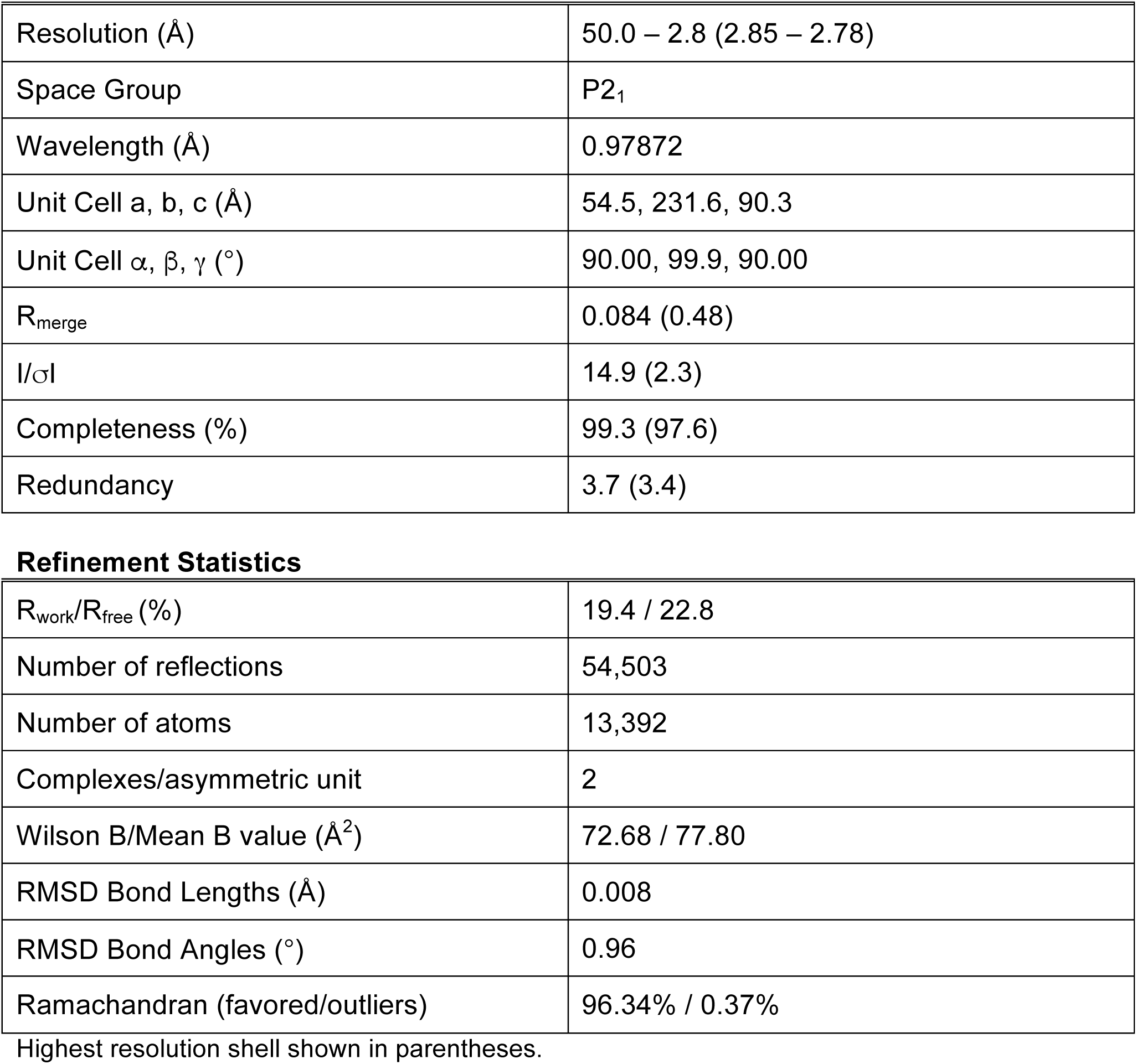
X-ray Data collection and refinement statistics.

|  |  |
| --- | --- |
| Resolution (Å) | 50.0 – 2.8 (2.85 – 2.78) |
| Space Group | P2<sub>1</sub> |
| Wavelength (Å) | 0.97872 |
| Unit Cell a, b, c (Å) | 54.5, 231.6, 90.3 |
| Unit Cell $\alpha$, $\beta$, $\gamma$ (°) | 90.00, 99.9, 90.00 |
| R<sub>merge</sub> | 0.084 (0.48) |
| I/σI | 14.9 (2.3) |
| Completeness (%) | 99.3 (97.6) |
| Redundancy | 3.7 (3.4) |
| **Refinement Statistics** |  |
| R<sub>work</sub>/R<sub>free</sub> (%) | 19.4 / 22.8 |
| Number of reflections | 54,503 |
| Number of atoms | 13,392 |
| Complexes/asymmetric unit | 2 |
| Wilson B/Mean B value (Å<sup>2</sup>) | 72.68 / 77.80 |
| RMSD Bond Lengths (Å) | 0.008 |
| RMSD Bond Angles (°) | 0.96 |
| Ramachandran (favored/outliers) | 96.34% / 0.37% |

Highest resolution shell shown in parentheses.

There are two RBPJ-MBP-SHARP/DNA complexes in the asymmetric unit (AU) of the crystals (Figure S1A) that overall are very structurally similar (RMSD 2.28 for 819 cα atoms), with even higher structural correspondence when either MBP (RMSD 0.07 for 297 cα atoms) or RBPJ (RMSD 0.66 for 388 cα atoms) are aligned individually (Figure S1C-E). The largest structural difference between the two complexes in the AU is a poorly ordered linker region in SHARP (Figure S1F), whereby the different conformations are likely influenced by different environments within the crystals. For clarity in subsequent figures, we do not show the MBP moiety and all structural comparisons are performed with chains G & H of RBPJ and SHARP, respectively, as these proteins chains have overall lower temperature factors.

### Bipartite interaction of SHARP with RBPJ

As shown in Figure 2A-B, SHARP interacts with two distinct surfaces on RBPJ, contacting the CTD and the BTD of RBPJ, which is consistent with previous binding studies^27^. Starting at the N-terminus of its RBPID, SHARP forms a β-hairpin motif that binds between the two β–sheets that compose the Ig domain of the CTD. The second β-strand of SHARP, residues 2788–2794, pairs with β-strand βg of the CTD, extending this from a three- to five-stranded β-sheet (Figure 2C). The binding mode of SHARP is structurally analogous to how the *Drosophila melanogaster* corepressor Hairless interacts with the CTD of Su(H) *(the fly CSL* ortholog)^18^, which is discussed further below. The SHARP/CTD interaction is followed by a short, approximately six residues, linker region that is poorly ordered and makes no contacts with RBPJ. The C-terminal portion of the SHARP RBPID binds in an extended fashion across the BTD in a manner that is structurally similar to the RAM domain of NICD^15,29,30^, and the corepressors FHL1 and RITA1 (Figure 2A-B and described below)^13,19^. The complex between SHARP and RBPJ is largely driven by hydrophobic interactions between nonpolar side chains on SHARP, and the CTD and BTD of RBPJ (Figure 2B); with SHARP residue L2791, as well as V2789 and Y2793, anchoring its interaction with the CTD; and I2811 anchoring the interaction between SHARP and the BTD. Additionally, there are key ionic interactions that appear to play auxiliary roles in complex formation, including salt bridges between E2786 of SHARP and R438 of the CTD (Figure 2C), and K2807 of SHARP and the BTD residue E259 (Figure 2B). It should also be mentioned that when complexed with SHARP, RBPJ maintains similar nonspecific and specific contacts with DNA, suggesting that SHARP binding does not affect the affinity of RBPJ for DNA, which is also consistent with previous binding studies^27^.

**Figure 2.**
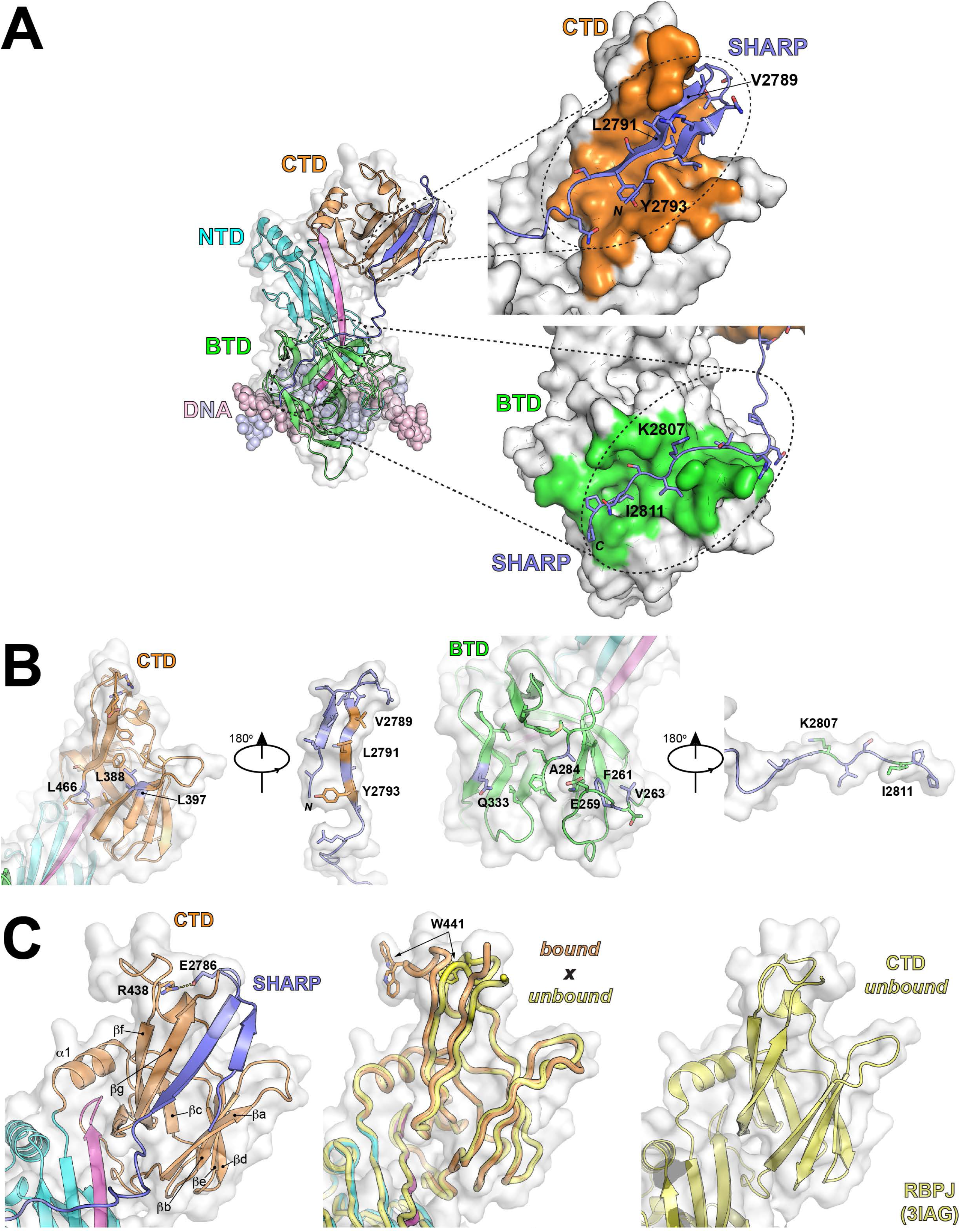
<u>X-ray structure of the RBPJ-SHARP corepressor complex bound to DNA</u>. (A) Ribbon diagram of RBPJ-SHARP-DNA complex with the NTD, BTD, and CTD of RBPJ colored cyan, green, and orange, respectively. SHARP is colored purple, and the DNA is colored light pink and light blue. Magnified views of the interaction of SHARP with the CTD (top) and BTD (bottom) of RBPJ. RBPJ is represented as a molecular surface with CTD and BTD residues that contact SHARP colored orange and green, respectively. The ribbon diagram of SHARP is colored purple with the side chains of SHARP that contact either the BTD or CTD shown as sticks. SHARP residues that were mutated and tested for activity in this study are labeled. (B) Open book representation of RBPJ-SHARP interfaces. (Left) Ribbon diagram with transparent surface of the CTD-SHARP interface, and (right), ribbon diagram with transparent surface of the BTD-SHARP interface. Side chains in RBPJ and SHARP that contribute to the interface are shown. Key residues at the interface that were mutated and tested for activity in this study are labeled. (C) Conformational changes in the CTD of RBPJ as a result of SHARP binding. (Left) Magnified view of the CTD-SHARP complex with labeled secondary structural elements of the CTD. The R438-E2786 salt bridge is also shown. (Middle) Structural overlay of RBPJ (bound) from RBPJ-SHARP complex with an unbound structure of RBPJ (PDBID: 3IAG). The RBPJ bound and unbound structures are colored orange and yellow, respectively. RBPJ residue W441, which undergoes a large conformational change in the bound structure to accommodate the R438-E2786 salt bridge, is shown in both structures. (Right) Ribbon diagram of the unbound RBPJ structure 3IAG.

As shown in Figure 2C, SHARP binding to RBPJ induces a structural change in CTD that results in translation of β-strands βf and βg outward, modestly expanding the CTD by as much as ~4 Å. A significant structural rearrangement is also observed for the loop that connects strands βe with βf, which strikingly repositions W441 from a buried to a solvent exposed conformation when comparing unbound RBPJ with the RBPJ/SHARP complex (Figure 2C). The repositioning of W441 allows for E2786 of SHARP to form a salt bridge with R438 of RBPJ. This structural rearrangement has not been observed for any other CSL structure determined to date.

### Structural comparison of coregulator binding sites on RBPJ

The intracellular domain of Notch (NICD) together with Mastermind (MAM) and RBPJ form a transcriptionally active ternary complex^6^. NICD interacts with high affinity to RBPJ^30–33^, binding the BTD and CTD of RBPJ via its RAM and ANK domains, respectively^15,16^. Figure 3A compares the interfaces that SHARP and NICD/MAM use to interact with RBPJ, illustrating the overlap of these binding sites. There is partial overlap of the SHARP and ANK/MAM binding sites on the CTD of RBPJ, whereas there is completely overlapping binding of SHARP and the RAM domain of NICD on the BTD of RBPJ. While SHARP and ANK/MAM use different strategies and target different key residues to interact with the CTD, clearly binding of SHARP and ANK/MAM to the CTD is mutually exclusive (Figure 3A). Despite the absence of sequence similarity between SHARP and the RAM domain of NICD, unexpectedly, SHARP interacts with the BTD in a manner that is structurally virtually identical to how RAM, as well as the corepressors FHL1 and RITA1, interacts with BTD (Figure 3B). As shown in Figure 3B, RAM and other coregulators that bind BTD have a characteristic hydrophobic tetrapeptide sequence (-ϕWϕP-) that is essential for binding BTD and serves as the linchpin for high affinity interactions. SHARP is lacking this and other conserved elements that contribute to interactions with BTD^31^. However, a structural alignment of SHARP with RAM and other BTD-binders reveals some sequence conservation of hydrophobic residues that play important roles in SHARP-RBPJ complex formation, including I2804, A2806, I2808, I2811, and P2812 (Figure 3B).

**Figure 3.**
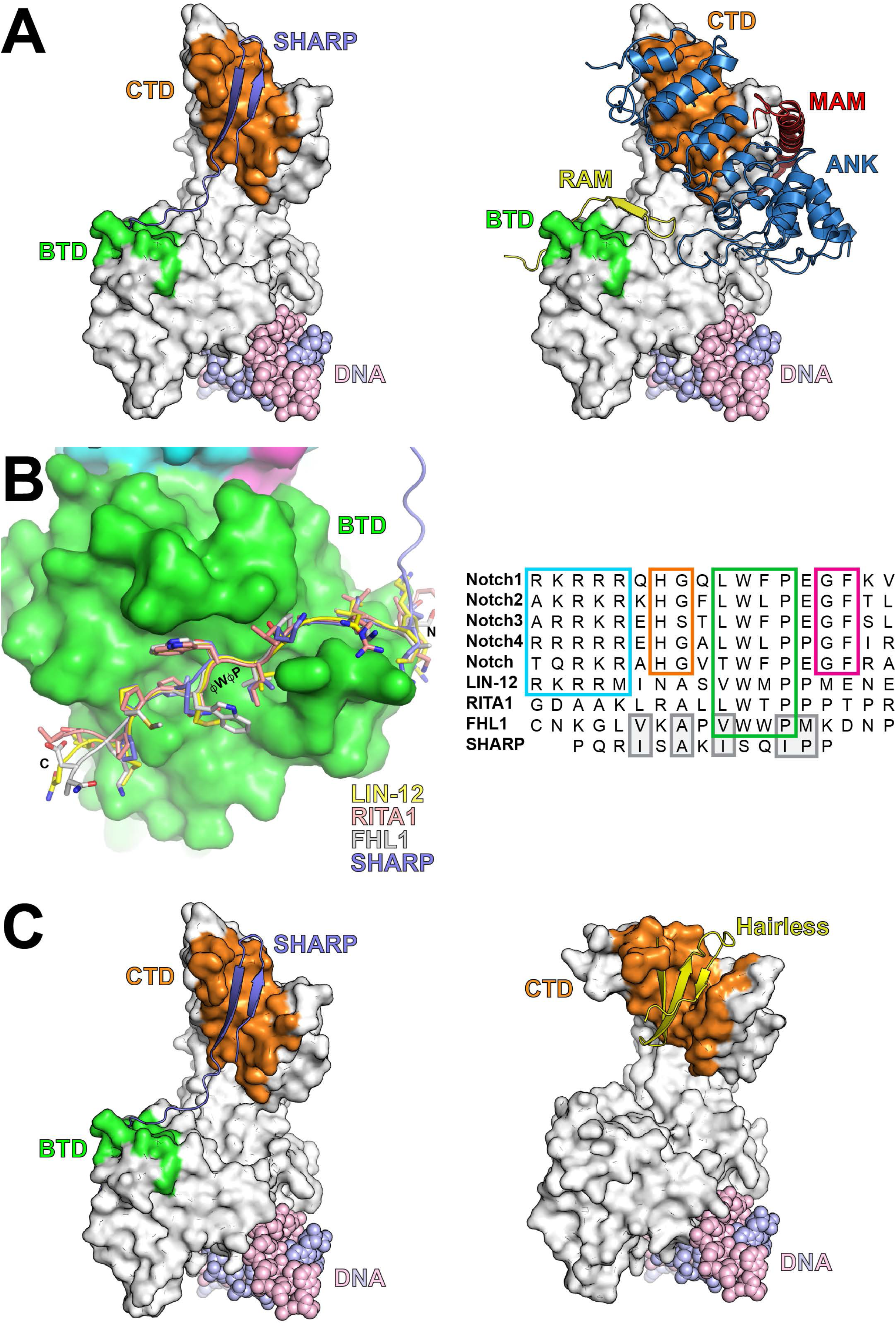
<u>Comparison of coregulator binding sites on RBPJ</u>. (A) Both SHARP and NICD bind the BTD and CTD of RBPJ. (Left) RBPJ/SHARP/DNA complex structure with RBPJ represented as a gray molecular surface, SHARP as a ribbon diagram colored purple, and the DNA as CPK colored light blue and pink. The RBPJ residues that contact SHARP in the BTD and CTD are colored green and orange, respectively. (Right) LAG-1/NICD/MAM/DNA complex structure (PDBID: 2FO1) with LAG-1 (C. *elegans* CSL ortholog) represented as a gray molecular surface. NICD is represented as a ribbon diagram with its RAM and ANK domains colored yellow and blue, respectively. MAM is depicted as a ribbon diagram and colored red. The DNA is represented in CPK and colored light blue and pink. RBPJ residues that contact RAM in the BTD are colored green and residues that contact ANK in the CTD are colored orange. (B) Structural alignment of SHARP and other coregulators that bind the BTD of CSL. (Left) The BTD of RBPJ is represented as a green molecular surface. SHARP, the RAM domain from LIN-12, RITA1 and FHL1 are colored purple, yellow, light pink, and gray, respectively. The hydrophobic tetrapeptide (ϕWϕP) is labeled. (Right) Sequence alignment of coregulators that bind the BTD of CSL, including the RAM domains of Notch from mammals, *D. melanogaster*, and *C. elegans*, and the corepressors RITA1, FHL1, and SHARP. The conserved ϕWϕP, which is absent in SHARP, is boxed in green. Other conserved regions in RAM that contribute to BTD binding are highlighted, including the basic region (blue), the -HG-motif (orange), and the -GF-motif (magenta). Structurally similar residues in SHARP that align with other BTD binders are boxed and highlighted in gray. (C) Structural similarity of RBPJ-SHARP and Su(H)-Hairless corepressor complexes. Despite no sequence similarity, both SHARP and Hairless bind the CTD in a structurally analogous manner. RBPJ and Su(H) are represented as a gray molecular surface with the SHARP and Hairless binding sites colored orange. SHARP and Hairless are shown as ribbon diagrams and colored purple and yellow, respectively. The DNA is shown as CPK spheres and colored light blue and light pink.

The corepressor Hairless is the major antagonist of Notch signaling in *Drosophila* and directly binds the CTD of Su(H) (fly *CSL* ortholog) with high affinity in order to repress Notch target gene transcription in flies^18,34,35^. While there is no sequence similarity between SHARP and Hairless, these two corepressors have evolved to bind the same CTD interface on RBPJ and Su(H) (Figure 3C). However, there are major structural differences in how SHARP and Hairless form complexes with the CTD. In contrast to SHARP, which only induces a modest conformational change in CTD, Hairless binding dramatically opens up the Ig domain of CTD, interacting with residues that form the hydrophobic core of CTD^18^. This allows Hairless to bind exclusively to the CTD with high affinity^34^.

The CTD is structurally similar to the RHR-C domains of transcription factors, such as NF-κB and NFAT^17^. Figure 4A shows the overlay of the CTD from RBPJ with the RHR-C domain from NFAT. Importantly, the CTD deviates from canonical RHR-C domains by the absence of β-strand a′ (βa′), which lies between the two β–sheetss that compose the Ig domain of a canonical RHR-C domain. Strikingly, SHARP binding to the CTD serves as a structural surrogate for βa′, occupying this region in the complex structure (Figure 4A), which has interesting implications for how binding sites have evolved on RBPJ and will be discussed further below (Figure 4B).

**Figure 4.**
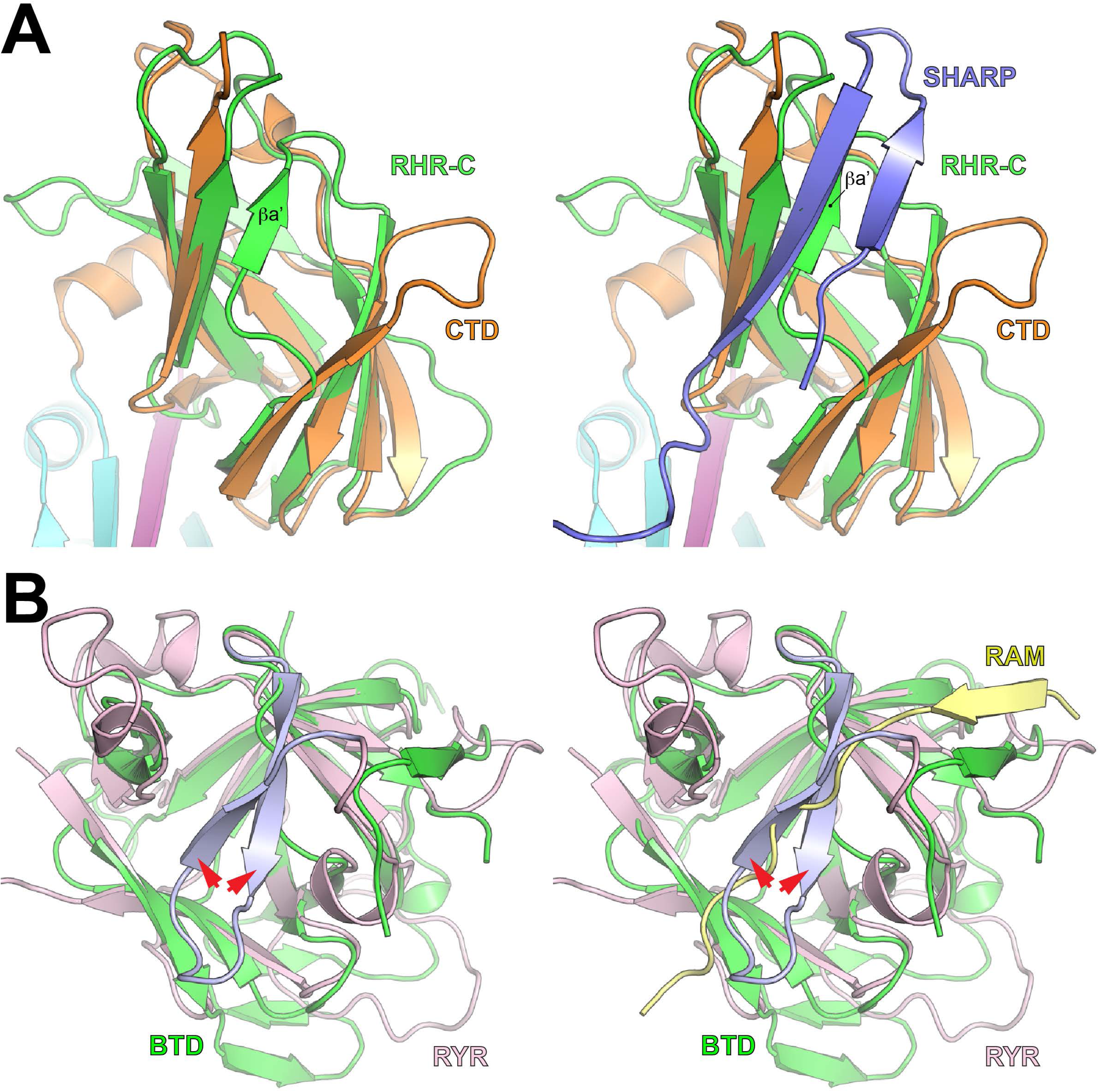
<u>Evolution of coregulator binding sites on CSL.</u>. (A) Structural comparison of the CTD of RBPJ with the RHR-C domain of NFAT. The ribbon diagram of the RBPJ-SHARP complex is colored similarly as in Figure 2, whereas NFAT is colored green. The NFAT β-strand βa’, which is absent in the CTD of RBPJ, but occupied by SHARP in the complex structure, is labeled. (B) Structural alignment of the BTD from RBPJ with a canonical β-trefoil fold from the ryanodine receptor (RYR). A canonical BTD is composed of 12 β-strands, in which four β-strands are arranged in a pseudo-threefold symmetrical arrangement. The atypical BTD of CSL is missing two of the 12 β-strands that compose a canonical BTD. The RAM domain of NICD acts as a structural surrogate, binding the region left absent by the missing β-strands. The BTD of CSL and RYR are colored green and light pink, respectively. The two β-strands that are missing in the CSL BTD fold are colored light blue and highlighted with red arrowheads. The RAM domain of NICD is colored yellow.

### Binding analysis of RBPJ and SHARP mutants

To gain further insights into RBPJ/SHARP complex formation, SHARP function, and to validate our structural studies, we used a combination of assays, both *in vitro* and in cells, to analyze structure-based mutants of RBPJ and SHARP. As shown in Figure 5 and Tables 2 and 3, we used isothermal titration calorimetry (ITC) to quantitate binding between SHARP and RBPJ mutants. SHARP binds RBPJ with high affinity (K_d_~5nM, Figure 5A and Table 2), wherein binding is enthalpically driven and entropically unfavorable, in accordance with the RBPID of SHARP being an intrinsically disordered region^27^. Consistent with its side chain burying ~84 Å^2^ in the SHARP-CTD complex, mutation of L2791 to alanine (L2791A) in SHARP resulted in greater than 350 fold decrease in binding (ΔΔG°=3.4kcal/mol, Figure 5C and Table 2). The SHARP mutants V2789A and Y2793A (Figure 5B,D and Table 2) also significantly reduced binding to RBPJ by >10 fold and >20 fold, respectively, in agreement with these residues also burying considerable amounts of surface area at the SHARP-CTD interface (V2789=54 Å^2^ and Y2793=118 Å^2^). The side chain of SHARP residue I2811 is buried at the BTD-SHARP interface, lying within a small hydrophobic pocket and burying a substantial 160 Å^2^. Mutation of I2811 to alanine (I2811A) results in more than a 60 fold reduction in binding (Figure 5F and Table 2). Mutation of K2807, which makes electrostatic interactions with E259 of the BTD, to alanine results in a much more modest effect on binding (~6 fold, Figure 5E and Table 2). Consistent with SHARP binding independently to the BTD and CTD, which results in an avidity effect^27^, single alanine mutants are unable to completely abrogate binding. However, the SHARP double mutant L2791A/I2811A, which targets mutations to key residues that interact with the BTD and CTD, results in a complete loss of binding in our ITC studies (Table 2).

**Figure 5.**
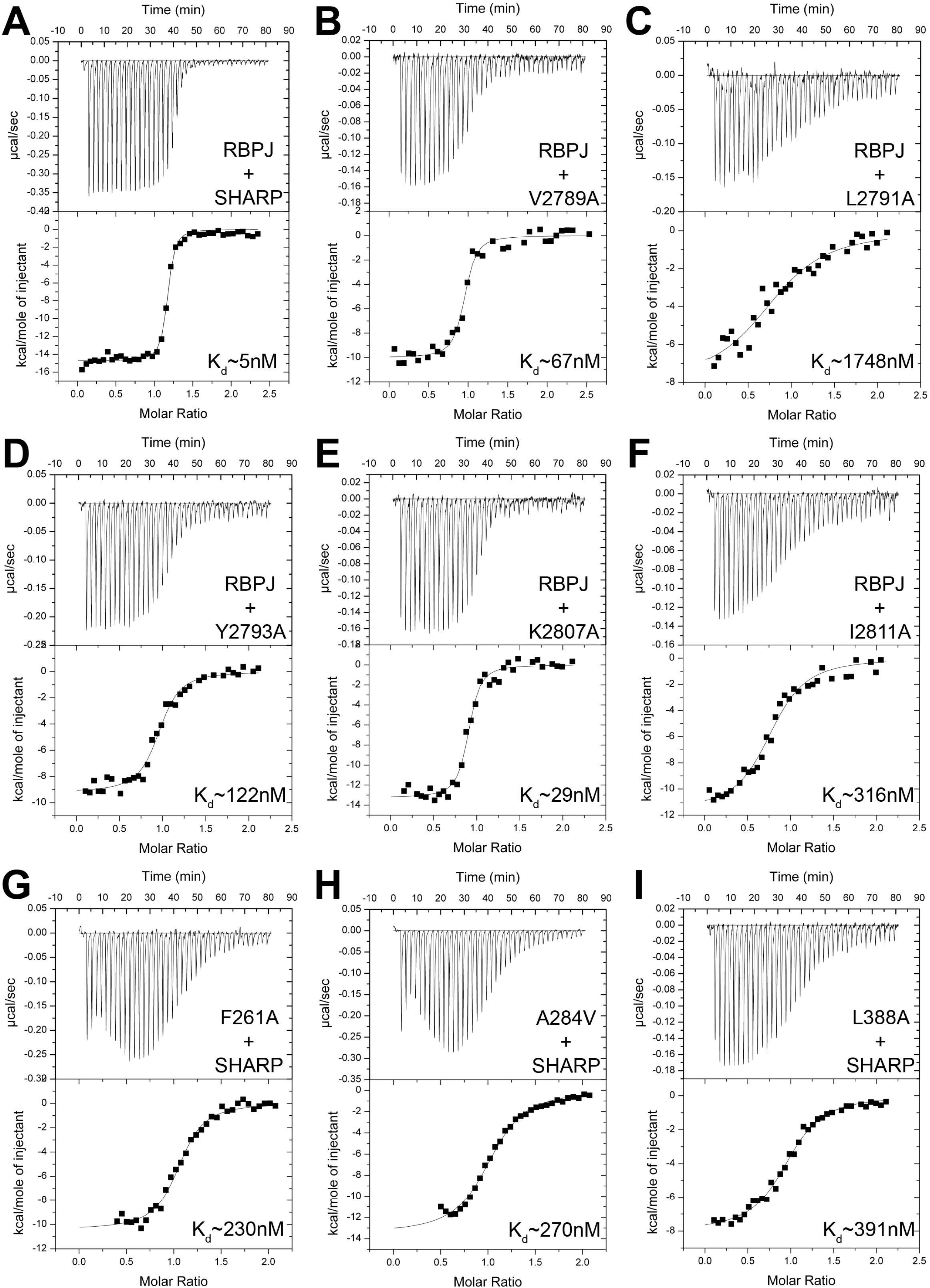
<u>ITC binding analysis of RBPJ and SHARP structure based mutants.</u>. Figure shows representative thermograms (raw heat signal and nonlinear least squares fit to the integrated data) for RBPJ binding to SHARP. Each experiment was performed at 25°C, with 40 titrations of 7 μl injections spaced 120 s apart. The dissociation constants (K_d_) shown for each experiment are indicated in Tables 2 and 3. (A) Native RBPJ and SHARP form a high affinity 1:1 complex with a 5 nM dissociation constant. The SHARP mutants L2791A (C), Y2793A (D), and I2811A (F) have the strongest effect on RBPJ-SHARP complex formation, whereas SHARP mutants V2789A (B) and K2807A (E) only modestly affect RBPJ binding. The RBPJ BTD mutants F261A (G) and A284V (H), and the CTD mutant L388A (I), strongly affect binding to SHARP.

**Table 2:** Calorimetric binding data for SHARP mutants and native RBPJ.

| SHARP | $K$ ($M^{-1}$) | $K_d$ ($\mu M$) | $\Delta G^\circ$ (kcal/mol) | $\Delta H^\circ$ (kcal/mol) | $-T\Delta S^\circ$ (kcal/mol) | $\Delta\Delta G^\circ$ (kcal/mol) |
| --- | --- | --- | --- | --- | --- | --- |
| WT | $2.0 \pm 0.5 \times 10^8$ | 0.005 | $-11.3 \pm 0.1$ | $-13.5 \pm 0.4$ | $2.2 \pm 0.3$ | -- |
| V2789A | $1.9 \pm 1.3 \times 10^7$ | 0.067 | $-9.9 \pm 0.4$ | $-11.0 \pm 0.9$ | $1.1 \pm 1.2$ | 1.4 |
| L2791A | $5.8 \pm 1.0 \times 10^5$ | 1.75 | $-7.9 \pm 0.1$ | $-7.8 \pm 0.8$ | $-0.1 \pm 0.7$ | 3.4 |
| Y2793A | $8.7 \pm 2.5 \times 10^6$ | 0.122 | $-9.4 \pm 0.2$ | $-9.0 \pm 0.2$ | $-0.4 \pm 0.4$ | 1.9 |
| K2807A | $3.6 \pm 0.8 \times 10^7$ | 0.029 | $-10.3 \pm 0.2$ | $-13.3 \pm 0.1$ | $3.0 \pm 0.1$ | 1.0 |
| I2811A | $3.3 \pm 0.6 \times 10^6$ | 0.316 | $-8.9 \pm 0.1$ | $-12.3 \pm 0.2$ | $3.4 \pm 0.2$ | 2.4 |
| L2791A/I2811A | NBD | --- | --- | --- | --- | --- |

All experiments were performed at 25°C. NBD represents no binding detected.

**Table 3:** Calorimetric binding data for native SHARP and RBPJ mutants.

| | RBPJ | $K$ ($M^{-1}$) | $K_d$<br>($\mu M$) | $\Delta G^\circ$<br>(kcal/mol) | $\Delta H^\circ$<br>(kcal/mol) | $-T\Delta S^\circ$<br>(kcal/mol) | $\Delta\Delta G^\circ$<br>(kcal/mol) |
| --- | --- | --- | --- | --- | --- | --- | --- |
| | WT | $2.0 \pm 0.5 \times 10^8$ | 0.005 | $-11.3 \pm 0.1$ | $-13.5 \pm 0.4$ | $2.2 \pm 0.3$ | -- |
| BTD<br>mutants | F261A | $4.5 \pm 0.9 \times 10^6$ | 0.23 | $-9.1 \pm 0.1$ | $-9.9 \pm 0.5$ | $0.8 \pm 0.4$ | 2.2 |
| | V263A | $3.2 \pm 0.4 \times 10^8$ | 0.003 | $-11.6 \pm 0.1$ | $-18.3 \pm 0.8$ | $6.7 \pm 0.9$ | -0.3 |
| | A284V | $4.0 \pm 1.0 \times 10^6$ | 0.27 | $-9.0 \pm 0.2$ | $-12.9 \pm 0.6$ | $3.9 \pm 0.7$ | 2.3 |
| | Q333A | $2.6 \pm 0.3 \times 10^8$ | 0.004 | $-11.4 \pm 0.1$ | $-16.8 \pm 0.7$ | $5.4 \pm 0.7$ | -0.1 |
| CTD<br>mutants | L386A | $7.5 \pm 3.5 \times 10^7$ | 0.016 | $-10.7 \pm 0.3$ | $-13.2 \pm 0.4$ | $2.5 \pm 0.7$ | 0.6 |
| | L388A | $2.6 \pm 0.5 \times 10^6$ | 0.391 | $-8.7 \pm 0.1$ | $-7.5 \pm 0.6$ | $-1.2 \pm 0.7$ | 2.6 |
| | L397A | $4.2 \pm 0.8 \times 10^7$ | 0.025 | $-10.4 \pm 0.1$ | $-11.3 \pm 0.2$ | $0.9 \pm 0.3$ | 0.9 |
| | L466A | $9.6 \pm 2.4 \times 10^7$ | 0.011 | $-10.9 \pm 0.1$ | $-13.3 \pm 0.2$ | $2.4 \pm 0.2$ | 0.4 |
| BTD/CTD | F261A/L388A | NBD | --- | --- | --- | --- | --- |

All experiments were performed at 25°C. NBD represents no binding detected.

Similarly, we performed ITC binding studies with structure-based RBPJ mutants and native SHARP. We performed differential scanning fluorimetry (DSF) to confirm that our RBPJ mutants are correctly folded (Figure S2A); EMSA to show that our RBPJ mutants bind DNA similarly to wild-type (WT) (Figure S2B); immunofluorescence microscopy to demonstrate that our RBPJ mutants properly localize to the nucleus (Figure S2C); and coimmunoprecipitation (CoIP) from cells to show that our RBPJ mutants still bind NICD (Figure S2D). However, we were unable to purify and test by ITC some RBPJ mutants that target key residues in CTD-SHARP complex, *e.g*. F468A, as these residues are buried within the hydrophobic core of the CTD and are required for proper folding of RBPJ. Nonetheless, as shown in Figure 5I and Table 3, the mutation of L388, which buries ~103 Å^2^ in the complex, to alanine results in an approximately 70 fold reduction in binding. Whereas the CTD mutants L386A, L397A, and L466A had only a minor effect on SHARP binding (Table 3), consistent with these residues burying much less surface area at the CTD-SHARP interface (L386=39 Å^2^, L397=3 Å^2^, and L466=44 Å^2^). Because SHARP binds the BTD of RBPJ in a structurally similar manner to the RAM domain of NICD, and the corepressors FHL1 and RITA1, we used a set of alanine mutants (F261A, V263A, A284V, and Q333A) that we have previously characterized for RAM, FHL1, L3MBTL3, and RITA1 binding to RBPJ (Figure 5G-H and Table 3)^12,13,19,36^. The BTD mutants F261A and A284V, which are more centrally located in the SHARP-BTD interface, have a stronger effect, significantly reducing binding by ~45 fold and ~50 fold, respectively (Figure 5G-H and Table 3). V263A and Q333A, which significantly reduced RAM binding to RBPJ^36^, but target more peripheral interactions in the BTD-SHARP complex, only modestly affect binding (Table 3). Interestingly, similar binding trends were observed when these BTD mutants were previously tested for interactions with FHL1 and RITA1^13,19^.

We also tested our SHARP mutants, and a subset of our RBPJ mutants, in cells using CoIP of exogenously expressed RBPJ and SHARP RBPID proteins, and a mammalian two-hybrid assay (Figure 6). Overall, we observed excellent correspondence between our ITC binding studies and our cellular assays. As shown in Figure 6A, the CoIP of SHARP mutants L2791A and I2811A with RBPJ is greatly reduced compared to wild-type SHARP, and the CoIP of the double mutant L2791A/I2811A is scarcely detectable on the Western blot (Figure 6A, lane 9); whereas, the CoIP of SHARP mutants V2789A, Y2793A, K2807A with RBPJ is only modestly affected (Figure 6A, lane 4–6), consistent with our ITC binding studies and the RBPJ-SHARP structure. Similarly, the CoIP of the BTD mutant F261A and the CTD mutant L388A with the RBPID of SHARP are greatly reduced compared to wild-type (Figure 6B, lanes 4–5), but the double mutant F261A/L388A was unable to CoIP SHARP (Figure 6B, lane 6). In our mammalian two-hybrid assays (Figure 6C-E), in which RBPJ and SHARP are fused to VP16 and Gal4, respectively, SHARP mutants L2791A and I2811A have the strongest effect on reporter expression, whereas V2798A, Y2793A, and K2807A only modestly affect reporter expression (Figure 6C). As expected from the complex structure, the SHARP double mutant L279A/I2811A, which does not bind RBPJ in our ITC assays, displayed no more than 1% expression from the reporter compared to wild-type (Figure 6C). The RBPJ mutants F261A and L388A were severely compromised for reporter activity (~1%), whereas no reporter activity was observed for the F261A/L388A double mutant. As a control, all RBPJ mutants fused to the VP16 transactivation domain activated a luciferase reporter containing 12 CSL binding sites (Figure 6E) and bound DNA in an EMSA (Figure S2B) similar to wild-type.

**Figure 6.**
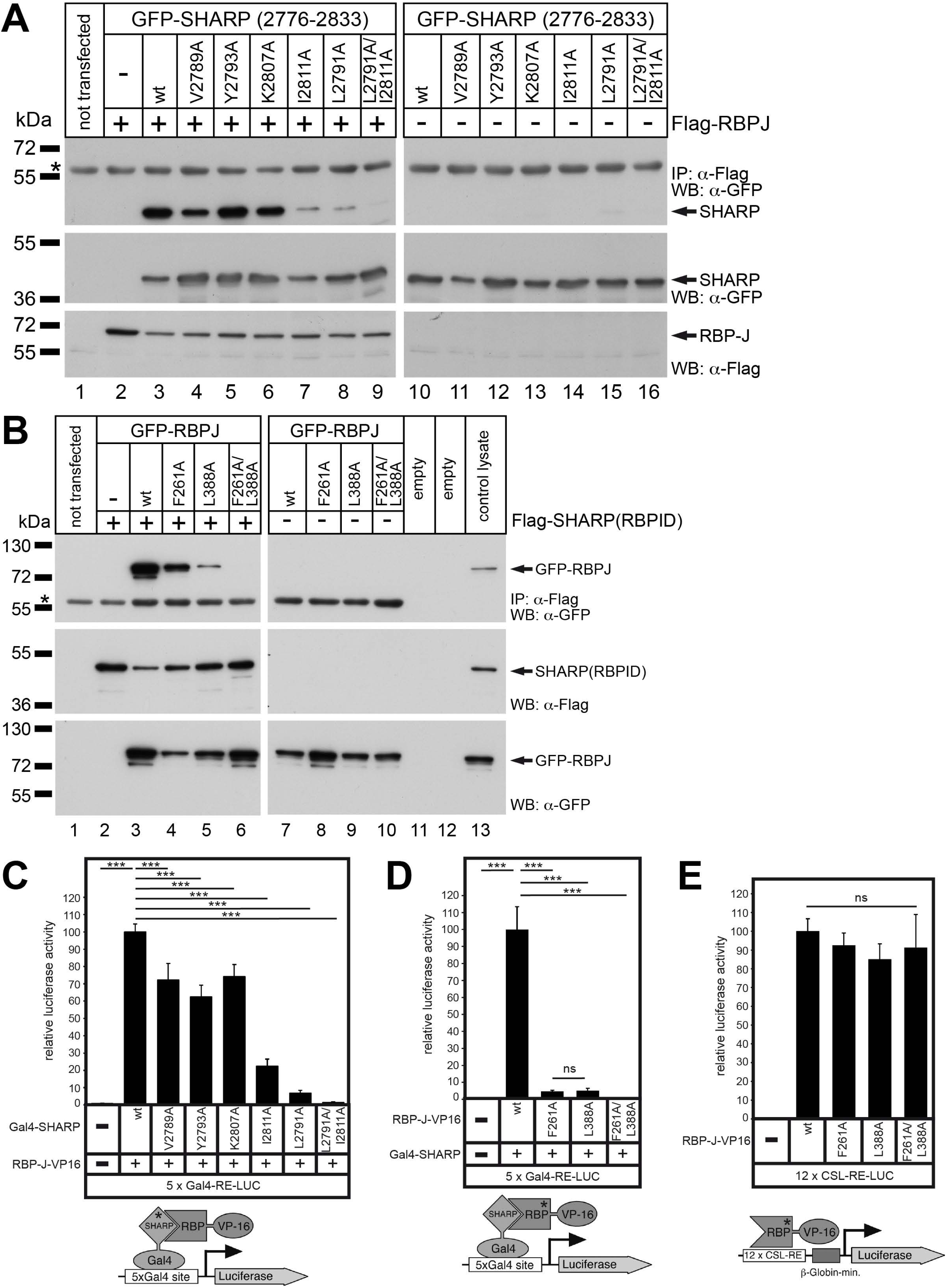
<u>Characterization of RBPJ and SHARP structure based mutants in cells.</u>. (A) Coimmunoprecipitation (CoIP) of RBPJ with wild-type and mutant SHARP constructs. HEK293 cells were cotransfected with GFP-SHARP (2766–2833), which corresponds to the RBPJ-interaction-domain of SHARP (RBPID), and Flag-RBPJ. Expression of GFP-SHARP (middle panel) and Flag-RBPJ (bottom panel) proteins was verified by Western blotting. CoIPs were performed 24h after transfection. The asterisk denotes the heavy chain of anti-Flag antibody used for immunoprecipitation. (B) CoIP of SHARP with wild-type and mutant RBPJ constructs. HEK293 cells were cotransfected with GFP-RBPJ and Flag-SHARP (RBPID). Expression of Flag-SHARP (middle panel) and GFP-RBPJ (bottom panel) proteins was verified by Western blotting. CoIPs were performed 24h after transfection. The asterisk denotes the heavy chain of anti-Flag antibody used for immunoprecipitation. (C-D) Analysis of structure based SHARP and RBPJ mutants using a mammalian two-hybrid assay. HeLa cells were cotransfected with the indicated Gal4-SHARP (300ng) and RBPJ-VP16 (150ng) constructs together with the pFR-Luc reporter (500ng), which contains five Gal4 DNA binding sites upstream of the luciferase gene. Relative luciferase activity was determined after cotransfection of the pFR-Luc reporter construct alone. Mean values and standard deviations are from at least six independent experiments. (E) Control experiments showing RBPJ-VP16 mutants are competent to bind and activate reporter similar to wild-type. HeLa cells were cotransfected with RBPJ-VP16 constructs (150ng) and the 12xCSL-RE luciferase reporter (500ng), which contains 12 CSL DNA binding sites upstream of the luciferase gene. Relative luciferase activity was determined after cotransfection of the reporter construct alone. Mean values and standard deviations are from four independent experiments [***P < 0.001, (ns) not significant, unpaired student's *t*-test].

### RBPJ/SHARP interaction is required to repress Notch target genes

In order to investigate the contribution of the RBPJ/SHARP interaction in the regulation of Notch target genes in cells, we analyzed the Notch target genes *Hes1* and *Hey1* in a mature T-cell line (MT) that lacks Notch activity. First, to interfere with RBPJ/SHARP-mediated repression in MT cells, we expressed the RBPID of SHARP as a GFP fusion protein either as wild-type (GFP-SHARP/RBPID^WT^) or the RBPJ-interacting defective L2791A/I2811A mutant (GFP-SHARP/RBPID^LI/AA^) (Figure 7A). Expression of GFP-SHARP/RBPID^WT^ leads to the upregulation, i.e. derepression, of *Hes1* and *Hey1*, whereas GFP-SHARP/RBPID^LI/AA^ has little to no effect on *Hes1 and Hey1* expression (Figure 7A). This suggests that GFP-SHARP/RBPID^WT^, but not GFP-SHARP/RBPID^LI/AA^, effectively outcompetes endogenous SHARP for binding to RBPJ. Next, we used CRISPR/Cas9 technology to deplete RBPJ (Figure 7B). Consistent with RBPJ repressor function, we observed robust upregulation of *Hes1* and *Hey1* in the absence of RBPJ (Figure 7B). Correspondingly, shRNA-mediated knockdown of RBPJ also resulted in the upregulation of *Hes1* and *Hey1* (Figure 7C). Importantly, reintroduction of wild-type RBPJ (RBPJ^WT^) in the CRISPR/Cas9-mediated RBPJ depleted background rescues the repression of *Hes1* and *Hey1* Notch target gene expression, whereas the SHARP-interacting defective RBPJ mutant F261A/L388A (RBPJ^FL/AA^) does not (Figure 7D). Altogether, our data demonstrates that the RBPJ/SHARP interaction is strongly required to repress transcription of Notch target genes in cells.

**Figure 7.**
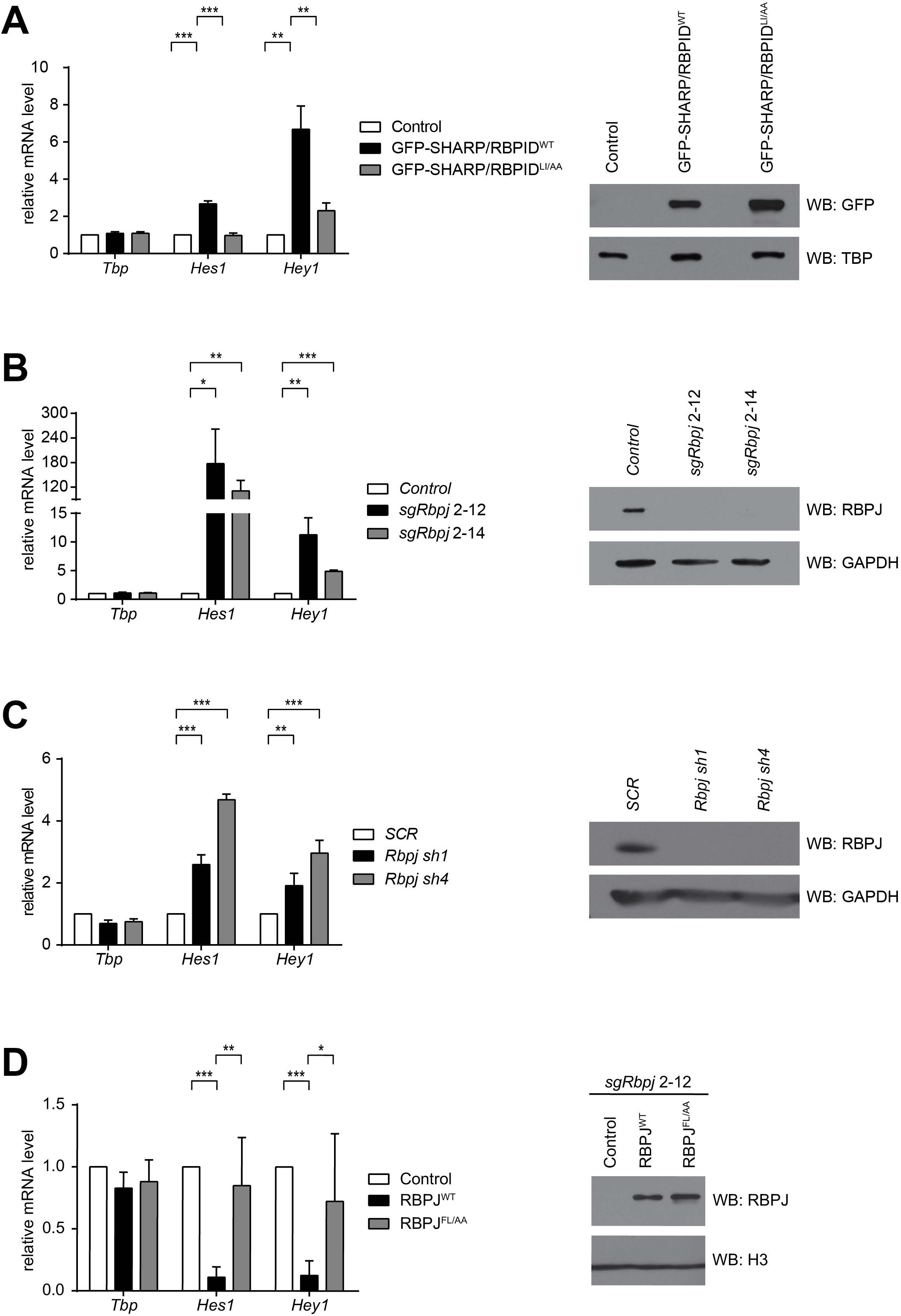
<u>The RBPJ-SHARP interaction is required for repression of Notch target genes in cells.</u>. (A) The wild-type SHARP RBPID (RBPID^WT^), but not the RBPJ-interacting defective SHARP RBPID (RBPID^LI/AA^) mutant, causes upregulation of Notch target genes in mouse mature T (MT) cells by outcompeting endogenous SHARP for RBPJ binding. MT cells were infected with plasmids encoding GFP-tagged SHARP (2776–2833), either wild-type (GFP-SHARP/RBPID^WT^, black bars) or the RBPJ-interacting defective mutant L2791A/I2811A (GFP-SHARP/RBPID^LI/AA^, gray bars), or an empty vector control (Control, white bars). (Left) Total RNA from MT cells infected with plasmids encoding GFP-SHARP/RBPID^WT^, GFP-SHARP/RBPID^LI/AA^, or an empty vector control, were reverse transcribed into cDNA and analyzed by qPCR using primers specific for *Tbp, Hes1* or *Hey1*. Data were normalized to the housekeeping gene *GusB* (glucuronidase β). Data shown represent the mean ± SD of triplicate experiments ([**] P < 0.01, [***] P < 0.001, unpaired Student’s *t*-test). (Right) Nuclear extracts (NE) were prepared from MT cells infected with GFP-SHARP/RBPID^WT^, GFP-SHARP/RBPID^LI/AA^, or an empty vector control and analyzed by Western blotting using a GFP antibody and with TBP used as a loading control. (B) RBPJ is required to repress Notch target genes *Hes1* and *Hey1* in MT cells as revealed by CRISPR/Cas9 depletion of RBPJ. (Left) Total RNA from wild-type *(Control)* or RBPJ depleted (clones *sgRbpj 2–12* and *sgRbpj 2–14)* MT cells was reverse transcribed into cDNA and analyzed by qPCR using primers specific for *Tbp, Hes1* or *Hey1*. Data were normalized to the housekeeping gene *GusB (glucuronidase β)*. Shown is the mean ± SD of triplicate experiments ([*] P < 0.05, [**] P < 0.01, [***] P < 0.001, unpaired Student’s *t*-test). (Right) Whole cell extracts (WCE) were prepared from wild-type *(Control)* or RBPJ depleted (clones *sgRbpj 2–12* and *sgRbpj 2–14)* MT cells and analyzed by Western blotting using an anti-RBPJ antibody. GAPDH was used as loading control. (C) *Hes1* and *Hey1* Notch target genes are upregulated upon shRNA-mediated *Rbpj* knockdown, but not with the *SCR* control shRNA. (Left) Total RNA from MT cells infected with shRNAs targeting *Rbpj (Rbpj sh1* or *Rbpj sh4)* or scrambled shRNA control (SCR) was reverse transcribed into cDNA and analyzed by qPCR using primers specific for *Tbp, Hes1* or *Hey1*. Data were normalized to the housekeeping gene *GusB (glucuronidase β)*. Shown is the mean ± SD of quadruplicate experiments ([**] P < 0.01, [***] P < 0.001, unpaired Student’s t-test). (Right) WCE was prepared from MT cells infected with shRNAs targeting *Rbpj (Rbpj sh1* or *Rbpj sh4)* or a scrambled shRNA control *(SCR)* and analyzed by Western blotting using a RBPJ antibody. GAPDH was used as loading control. (D) Expression of wild-type RBPJ (RBPJ^WT^), but not the SHARP-interacting defective mutant F261A/L388A (RBPJ^FL/AA^), in the RBPJ-depleted background rescues the repression of Notch target genes. (Left) Total RNA from *sgRbpj 2–12* MT cells infected with empty vector (Control), RBPJ^WT^ or RBPJ^FL/AA^ was reverse transcribed into cDNA and analyzed by qPCR using primers specific for *Tbp, Hes1* or *Hey1*. Data were normalized to the housekeeping gene *GusB (glucuronidase β)*. Shown is the mean ± SD of three independent experiments measured twice each ([*] P < 0.05, [**] P < 0.01, [***] P < 0.001, unpaired Student’s t-test). (Right) NEs were prepared from *sgRbpj 2–12* MT cells infected with empty vector (Control), RBPJ^WT^ or RBPJ^FL/AA^ and analyzed by Western blotting using anti-RBPJ antibodies. Histone H3 was used as a loading control.

## Discussion

Notch is a highly conserved signaling pathway that ultimately results in changes in gene expression, a process that is controlled by the transcription factor CSL (Figure 1A)^1,2^. CSL forms an activation complex with the intracellular domain of the Notch receptor (NICD) and the coactivator Mastermind (MAM) to activate transcription from Notch target genes (Figure 1B)^15,16^. This activity of CSL is required at all Notch target genes and is conserved in all metazoans. CSL can also function as a transcriptional repressor, by interacting with corepressors such as SHARP^8,9,27^, L3MBTL3^12^, FHL1^11,19^, and RITA1^13,14^ in mammals, and Hairless in *Drosophila^18,34,35^*; however, its role as a repressor, in particular in mammals, is not well understood. In order to begin to address this gap in our understanding, here we determine the X-ray structure of the RBPJ-SHARP corepressor complex bound to DNA and use a multitude of *in vitro* and cellular assays to characterize structure based RBPJ and SHARP mutants in order to better understand CSL-corepressor function.

Together with previous studies, the complex structure provides a detailed mechanism for how the corepressor SHARP interacts with the transcription factor RBPJ to repress transcription. Prior to binding RBPJ, the RBPID of SHARP is intrinsically disordered^27^, which then forms a bipartite interaction with RBPJ. As shown in Figure 2, the N-terminal portion of the RBPID assumes a β-hairpin motif that interacts with the CTD of RBPJ, resulting in a modest conformational change in the CTD; the SHARP-CTD interaction is followed by a poorly structured linker and then an extended region that binds across a nonpolar surface on the BTD. Thus, SHARP forms a high affinity complex with RBPJ by interacting with two distant binding surfaces on RBPJ, which individually are low affinity, but when tethered by the linker region result in an avidity effect^27^. RBPJ-SHARP complex formation localizes the RRM and SPOC domains of SHARP to Notch target genes (Figure 1C). In the case of SPOC, this recruits other factors involved in transcriptional repression, such as NCoR/SMRT^20,24^, CtIP (C-terminal binding protein interacting protein)/CtBP (C-terminal binding protein)^23^, and ETO (eight-twenty-one)^22^, as well as the KMT2D coactivator complex^24^. Whether the RRM domains of SHARP recruit Xist lncRNA or other lncRNAs, such as SRA (steroid receptor coactivator)^37^, to Notch target genes remains to be determined

Analogous to SHARP, NICD also binds the BTD and CTD of RBPJ (Figure 1B and 3A); however, the two complexes with RBPJ are mutually exclusive (Figure 3A) and the distribution of binding energies for surfaces on RBPJ is distinct. In mammals, the RAM domain of NICD forms a high affinity interaction with the BTD^30–33^, comparable to the affinity of SHARP for RBPJ^27^, and is structurally very similar to how SHARP interacts with the BTD despite no sequence similarity between the two proteins (Figure 3B). The ANK domain of NICD, however, binds the CTD of RBPJ in a manner that is structurally very different than SHARP and the ANK-CTD interaction is extremely weak, barely discernible in some binding assays^27,30,32,33^. MAM is then required to stabilize the RBPJ-NICD-MAM ternary complex. The functional consequences of these structural and binding differences between SHARP and NICD are currently unclear, but it may be important for the dynamic association/dissociation of RBPJ-coregulator complexes and the lifetimes of these complexes in the nucleus. Certainly, single molecule studies of wildtype and mutant SHARP molecules will be important for addressing the dynamics of RBPJ-coregulator complexes in live mammalian cells.

Our cellular studies clearly demonstrate that CRISPR/Cas9-mediated depletion of RBPJ in MT cells results in derepression of the well-established Notch target genes *Hes1 and Hey1* (Figure 7)^38^. Moreover, reconstitution of repression by wild-type RBPJ, but not a SHARP-binding mutant, strongly suggests that SHARP is the primary corepressor that mediates repression via RBPJ at Notch target genes in lymphocytes, and likely other cells and tissues, which is consistent with previous *in vivo* studies of SHARP in mice^8,39,40^. These data are also consistent with research in other experimental systems that show that loss of CSL results in transcriptional derepression at some, but not all, Notch target genes^41,42^. Why RBPJ-SHARP corepressor complexes are recruited to some Notch targets, but not others, remains an open question. Clearly, the use of RBPJ mutants defined here that affect SHARP interactions, but leave NICD interactions largely intact, will be instrumental in addressing this and other questions regarding the function of RBPJ as a transcriptional repressor.

Classical models of transcriptional regulation in the Notch pathway posit that CSL is constitutively bound to DNA, and that corepressors are replaced by NICD when signaling is activated^43,44^. Given the comparable high affinities of both SHARP and NICD for RBPJ^27,30^, as well as the more modest affinity of CSL for DNA (~0.5–1.0 μM)^45,46^, it is difficult to reconcile how cofactor exchange on DNA, *i.e*. NICD replacing SHARP on RBPJ, might work at the molecular level. The affinities rather suggest that preformed RBPJ-corepressor complexes and RBPJ-NICD-MAM coactivator complexes are dynamically associating/dissociating with sites on the DNA. This scenario is supported by recent data by Bray and colleagues investigating the DNA-binding dynamics of Su(H) complexes with NICD and Hairless in Notch-ON and Notch-OFF states in the salivary gland of *Drosophila melanogaster^47^*.

In contrast to the CSL-NICD-MAM activator complex, CSL-corepressor interactions appear to be less conserved across disparate organisms. For example, while SHARP, also known as SPEN in *Drosophila* and DIN-1 in *C. elegans*, is conserved from nematodes to flies to mammals^48^, the RBPID of SHARP is only conserved in vertebrates^27^. Similarly, the corepressor Hairless, which is the major antagonist of Notch signaling in *Drosophila*, is not conserved outside of insects and crustaceans^49^. However, while the corepressors that bind CSL are not strictly conserved across disparate organisms, interestingly, the corepressor binding sites on CSL are conserved. For example, SHARP, as well as the corepressors RITA1, FHL1, and likely L3MBTL3, binds to the BTD of RBPJ in a structurally similar manner (Figure 3B); and strikingly, despite no sequence conservation between SHARP and Hairless, both proteins bind to the same surface of the CTD of CSL (Figure 3C). Thus, it seems likely that other species-specific transcriptional corepressors that bind CSL will be identified, but they will likely utilize the aforementioned conserved binding surfaces on the BTD and CTD to interact with CSL.

In a broader context, CSL provides an interesting example for how ligand binding sites evolve in proteins. CSL is composed of two types of domains – the NTD and CTD are immunoglobulin domains that are structurally related to the rel homology region (RHR) of proteins, such as NF-κB and NFAT, and the BTD contains a β-trefoil fold that is structurally related to growth factors such as interleukins and FGF^17^. As shown in Figure 4A, the canonical RHR-C fold contains a β-strand, βa’, that lies in between β-strands βa and βg; however, the CTD of CSL is missing βa’ typically found in RHR-C domains. The ligand, in this case the corepressor SHARP, serves as a structural surrogate binding precisely in the region where the missing βa’ strand would lie. Strikingly, this is the second example in CSL proteins where this type of ligand evolution has occurred. The first example, as shown in Figure 4B, was uncovered following the structure determination of the CSL-NICD-MAM activation complex^15^, whereby it was shown that the BTD of CSL is atypical, as it deviates from the canonical β-trefoil fold such that it is missing two of the 12 strands of the consensus fold^17^. In this instance, the ligand, the RAM domain of NICD, again serves as a structural surrogate, binding across the BTD exactly where the two missing β-strands would normally lie.

Finally, because of its association with human disease, there have been wide-ranging efforts to pharmacologically target the Notch pathway, but the majority of this work has focused on reagents that blunt overactive Notch signaling, *e.g*. in T-cell acute lymphoblastic leukemia^4,5^. However, there are numerous human diseases, including certain types of cancer, cardiovascular defects, and congenital syndromes that are associated with insufficient Notch signaling^3^. Our structure-function studies of the RBPJ-SHARP corepressor complex provide a roadmap for developing reagents that lead to the derepression of Notch target genes, which have the potential to become new treatment options for diseases connected to deficiencies in Notch signaling.

## Methods

### Protein expression and purification

The cloning, expression, and purification of *Mus musculus* core RBPJ, residues 53–474, and SMT3-SHARP, residues 2776–2833, which corresponds to the RBPJ-interacting region, were described previously^27^. An MBP-SHARP fusion protein was used to crystallize the RBPJ/SHARP/DNA complex. SHARP, corresponding to residues 2776–2820, was cloned into pMALX-E, which encodes maltose binding protein with the following surface entropy reduction mutations D82A, K83A, E172A, N173A, and K239A to aide in crystallization. The MBP-SHARP fusion construct was overexpressed in Tuner *E. coli* and cells were lysed by sonication. The lysate was incubated with amylose resin and eluted with 10 mM maltose. The MBP-SHARP fusion protein was further purified by size exclusion chromatography.

### Crystallization and Data Collection

RBPJ/MBP-SHARP/DNA complexes were setup in a 1:1.1:1.1 molar ratio and screened for crystallization conditions using an Art Robbins Phoenix Crystallization Robot at 4°C. The RBPJ/MBP-SHARP/DNA complex crystallized in 100mM Bis-Tris pH 6.6, 100mM NaCl, 40% PEG 400, and 200mM NDSB-256. Crystals were harvested, flash frozen in liquid nitrogen, diffraction data were collected at the Advanced Photon Source, beamline 21-ID-F (LS-CAT). The RBPJ/MBP-SHARP/DNA crystals nominally diffract to 2.8Å and belong to the spacegroup P2_1_ with unit cell dimensions a=54.5 Å, b=231.6 Å, c=90.3 Å, and ß=99.88°.

### Structure Determination, Model Building, and Refinement

Molecular replacement with Phaser^50^ was used to determine the RBPJ/MBP-SHARP/DNA complex structure using the following search models DNA (3BRG), CSL (3IAG), and MBP (3OB4). Two RBPJ/MBP-SHARP/DNA complexes were identified in the asymmetric unit. Phenix was used for the initial stages of refinement^51^. Manual model building was performed with COOT^52^. TLS parameters were generated and the model was subsequently refined using BUSTER^53^ and validated with MolProbity^54^. The final RBPJ/MBP-SHARP/DNA model was refined to an R_work_ = 18% and R_free_ = 23% with good overall geometry (see Table 1). PyMOL (The PyMOL Molecular Graphics System, Version 2.0 Schrödinger, LLC) was used to generate figures and perform structural overlays. The PISA server was used to analyze protein interfaces^55^.

Coordinates and structure factors have been deposited into the Protein Data Bank (XYZ1)

### Isothermal titration calorimetry (ITC)

ITC experiments were performed using a Microcal VP-ITC micocalorimeter. For all binding reactions, SMT3-SHARP (2776–2883) at ~100 μM was placed in the syringe concentrations and RBPJ (53–474) at ~10 μM was placed in the cell. Titrations consisted of an initial 1μl injection followed by 39 7μl injections. ITC binding experiments were performed in 50 mM sodium phosphate pH 6.5, 150 mM NaCl at 25°C. Samples were buffer matched using size-exclusion chromatography or dialysis. The raw data were analyzed using ORIGIN and fit to a one site binding model.

### Cell culture and preparation of cell extracts

Mouse hybridoma mature T (MT) cell line was grown in Iscove’s Modified Dulbecco Medium (IMDM, Gibco) supplemented with 2% FCS, 0.3 mg/l peptone, 5 mg/l insulin, nonessential aminoacids and penicillin/streptomycin. Cells were grown at 37°C with 5% CO_2_. HeLa (ATCC: CCL2), HEK-293 (ATCC: CRL1573), 293T and *Phoenix™* packaging cells *(Orbigen*, Inc., San Diego, CA, USA) were cultivated in Dulbecco’s Modified Eagle Medium (DMEM, Gibco) supplemented with 10% fetal calf serum (FCS) and penicillin/streptomycin.

### DNA transfection

HEK-293 and HeLa cells were transfected using the Nanofectin transfection reagent (PAA) according to the manufacturer’s instructions.

### Coimmunoprecipitation experiments

HEK-293 cells were transfected with the indicated constructs for expression of GFP- and Flag-tagged wild-type and mutant proteins. 24 hours after transfection cells were lysed with 600 μl CHAPS lysis buffer [10 mM 3-[(3-Cholamidopropyl)dimethylammonio]-1-propanesulfonate hydrate (CHAPS, Merck), 50 mM Tris-HCl (pH 7.8), 150 mM NaCl, 5 mM NaF, 1 mM Dithiothreitol (DTT, Merck), 0.5 mM Phenylmethanesulfonyl fluoride (PMSF, Merck) and 40 μl/ml “Complete Mix” protease inhibitor cocktail (Roche)]. The extracts were incubated with 40 μl agarose-conjugated anti-Flag antibody (M2, Sigma) at 4°C overnight. Precipitates were washed 6 to 8 times with CHAPS lysis buffer and finally resuspended in SDS-polyacrylamide gel loading buffer. For Western blotting the proteins were resolved in SDS-polyacrylamide gels and transferred electrophoretically at room temperature to PVDF membranes (Millipore) for 1 h at 50 mA using a Tris-glycine buffer system. The membranes were pre-blocked for 1 h in a solution of 3% milk powder in PBS-T (0.1% Tween 20 in PBS) before adding antibodies. The following antibodies were used: anti-GFP (7.1/13.1, mouse monoclonal IgG, secondary antibody peroxidase conjugated sheep anti-mouse IgG, NA931V, GE healthcare) or anti-Flag (M5, Sigma; secondary antibody, NA931V, GE healthcare).

### In vitro protein translation

The *in vitro* protein translations were performed using the TNT-assay (#L4610) from Promega according to manufacturer's instructions. Prior to EMSAs the *in vitro* translations of RBPJ (wt) and mutant proteins were monitored by western blotting using an anti-Flag antibody (M5, Sigma).

### Electro Mobility Shift Assay (EMSA)

Reticulocyte lysates from *in vitro* translations were used for electromobility shift assays (EMSAs) in a binding buffer consisting of 10 mM Tris-HCl (pH 7.5), 100 mM NaCl, 0.1 mM EDTA, 0.5 mM DTT, and 4% glycerol. For binding reaction, 2 μg poly(dI-dC) (GE healthcare) and approximately 0.5 ng of ^32^P-labelled oligonucleotides were added. The sequence of the double-stranded oligonucleotide FO-233 (Supplementary Table S1) corresponds to the two RBP-J-binding sites within the EBV TP-1 promoter. Super shifting of complexes was achieved by adding 1 μg of anti-Flag (M5, Sigma) antibody. The reaction products were separated using 5% polyacrylamide gels with 1x Tris-glycine-EDTA at room temperature. Gels were dried and exposed to X-ray films (Kodak).

### Fluorescence microscopy

HeLa cells were cultured on glass cover slips in a 25-well plate (Bibby Sterilin Ltd) at a density of 10^5^ cells per cm^2^. After 16 h cells were transfected with 400 ng of expression plasmids using the Nanofectin transfection reagent (see above). 24 h after transfection cells were rinsed with PBS, fixed with 4% paraformaldehyde (PFA, Merck) in PBS (pH = 7.5). Specimens were embedded in “ProLong© Gold antifade” reagent (Invitrogen) supplemented with 2-(4-carbamimidoylphenyl)-1H-indol-6-carboximidamide (DAPI) and stored at 4 °C overnight. Pictures were taken using a fluorescence microscope (IX71, Olympus) equipped with a digital camera (C4742, Hamamatsu), and a 100-W mercury lamp (HBO 103W/2, Osram). The following filter sets were used: Green, (EGFP) ex: HQ470/40, em: HQ525/50, blue (DAPI) D360/50, em: D460/50.

### Luciferase assay

HeLa cells were seeded in 48-well plates at a density of 20 × 10^4^ cells. Transfection was performed with Nanofectin reagent (see above) using 1 μg of reporter plasmid alone or together with various amounts of expression plasmid (given in the corresponding figure legends). After 24 h luciferase activity was determined from at least six independent experiments with 20 μl of cleared lysate in an LB 9501 luminometer (Berthold) by using the luciferase assay system from Promega.

### Infection of hybridoma mature T-cell line

5 × 10^6^ *Phoenix™* cells were seeded and 24 h later they were transfected with the plasmid DNA of choice. Briefly, 20 μg of DNA were mixed with 860 μl of H_2_O and 120 μl of 2 M CaCl_2_ and mixed by vortexing. The DNA solution was transferred dropwise to 1 ml of 2 x HBS buffer (50 mM HEPES pH 7.05, 10 mM KCl, 12 mM Glucose, 280 mM NaCl, 1.5 mM NaHPO_4_) while vortexing and the solution was incubated 20 min at room temperature. In the meantime, 25 μM Chloroquine solution (Sigma-Aldrich) was added to the *Phoenix™* cells (1 μl/ml) and the cells were incubated for 10 min. The DNA solution was added to the cells and 12 h later the medium was replaced. After 24 h of incubation, the medium containing the retroviral suspension was filtered and 2 mg/ml Polybrene (Sigma-Aldrich) solution was added (1 μl/ml). Fresh medium was added to the *Phoenix™* cells that were maintained in culture for further infections. The retroviral solution was used for spin infection of MT cells by centrifuging 45 min at 1800 rpm at 37°C. In total, four spin infections were performed over two days. Positively infected cells were selected with puromycine (Serva) or blasticidin (Gibco) and, eventually, GFP positivity was analyzed using a BD FACS Calibur.

### *Generation of CRISPR/Cas9* depleted MT cells

CRISPR/Cas9 *Rbpj* depleted MT cells were generated as follows: 3 × 10^6^ 293T cells were seeded and, after 24 h, transfected with 2.5 μg psPAX, 1 μg pMD2G and 3.33 μg of the desired lentiCRISPR v2 vector using Lipofectamine 2000 Transfection Reagent (Invitrogen 11668–019) accordingly to manifacturer’s instructions. After at least 6 h of incubation at 37°C the medium was replaced with fresh one and 48 h post-transfection the supernatant was filtered, supplemented with polybrene and used for infection of MT cells. Positively infected cells were selected with puromycin and dilutions were performed to establish single cell clones. Individual clones were screened by Western blotting versus RBPJ.

### shRNA knockdown

For the knockdown in MT cells, the pLKO.1 TRC1 shRNA library (SIGMA-ALDRICH) was used. Transfection of 293T cells and infection and selection of MT cells was performed as previously described. Sequence of the hairpins used in this study is indicated in Table S1.

### Constructs

The expression plasmid pcDNA3-Flag-mNotch-1-IC (Flag-NICD) and the luciferase reporter construct pGa981/6 (12 x CSL-RE-LUC) were previously described^14^. The Gal4-reporter plasmid pFR-Luc (5 x Gal4-RE-LUC) was previously described^9^. The pMSCV-FLAG-hRBP-J IRES Blasticidin was kindly provided by Dr. R. Liefke. All oligonucleotides used in this study are listed in Table S1. PCR products were cloned in the pSC-A-amp/kan (Agilent Technologies 240205–5), digested with the selected restriction enzymes (New England Biolabs) and cloned into the destination vectors accordingly to Table S2. All plasmids were analyzed by sequencing. The pcDNA 3.1 Flag2 (Invitrogen) was commercially acquired while the pMY BioTip60 IRES-GFP was previously described.

An engineered CRISPR/Cas9 resistant mouse RBPJ cDNA was synthetized at GENEART/Life Technologies and inserted into the pcDNA3.1 Flag2 via NotI digestion. The RBPJ mutants R218H, F261A, L388A and the F261A/L388A double mutant was generated by site directed mutagenesis using the QuikChange II XL Site-Directed Mutagenesis Kit (Agilent Technologies 200521–5) accordingly to manufacturer’s instructions with primers listed in Table S1 and using the pcDNA3.1 Flag-mRBPJ wt CRISPR/Cas9 resistant plasmid as template. The mRBPJ wt and RBPJ F261A/L388A CRISPR/Cas9 resistant cDNAs were subcloned into the pMY-Bio-IRES Blasticidin. The mouse RBPJ-VP16 expression plasmids (pcDNA3.1-Flag-2-mRBPJ-VP16 wt, F261A, L388A and F261A/L388A) were constructed as follows: The stop codon was deleted by a mRBP-J specific PCR fragment (mRBP_VP16_UP, mRBP-TAA_DO) resulting in the pcDNA3.1-Flag-2-mRBPJΔstop constructs. A VP16 specific PCR-fragment (VP16_XhoI_UP, VP16_XbaI_DO) was inserted into the corresponding sites of pcDNA3.1Flag-2-mRBPJΔstop constructs resulting in the pcDNA3.1-Flag2-mRBPJ-VP16 plasmids. The Gal-4-Mint (2776–2833) and the EGFP-Mint (2776–2833) constructs were generated by PCR assisted cloning (Gal-Mint_F, Gal-Mint_R) into the BamHI/XbaI sites of pFa-CMV (Stratagene) and pEGFP-C1 (Clontech), respectively. The mutated constructs (V2785A, Y2793A, K2807A, I2811A, L2791A and I2811A/L2791A) were obtained by site directed mutagenesis.

The lentiCRISPR v2 was a gift from Dr. F. Zhang (Addgene plasmid # 52961). The CRISPR/Cas9 guides were designed using the online tool available at http://crispr.mit.edu/. The desired 5’ overhangs were added and oligos were phosphorylated, annealed and ligated into the lentiCRISPRv2 predigested with BsmBI.

### RNA extraction, RT-PCR and qPCR

Total RNA was purified using Trizol reagent (Ambion, 15596018) accordingly to manufacturer’s instructions. 1 μg of RNA was reverse transcribed in cDNA using random hexamers and M-MuLV reverse transcriptase (NEB). qPCRs were assembled with Absolute QPCR ROX Mix (Thermo Scientific, AB-1139), gene-specific oligonucleotides and double-dye probes (see Table S1) and analyzed using the StepOne Plus Real Time PCR system (Applied Biosystem). Data were normalized to the housekeeping gene *glucuronidase β (GusB)*.

### Preparation of protein extracts and Western Blotting from MT cells

Whole Cell Extract (WCE) was prepared as follows. Briefly, cells were washed twice in PBS, lysed in WCE buffer (20 mM Tris-HCl pH 8.0, 150 mM NaCl, 1% NP-40, 10% glycerol, 0.5 mM Na_3_VO_4_, 10 mM NaF, 1 mM PMSF, 1 x Protease inhibitor cocktail mix) and incubated 20 min on ice. After centrifuging 15 min at 13200 rpm at 4°C, the supernatant was recovered.

The Nuclear Extract (NE) from MT cells overexpressing the SHARP constructs was prepared as follows. Briefly, cells were washed with PBS and resuspended in Buffer A (20 mM Hepes pH 7.9 / 20 mM NaCl / 5 mM MgCl_2_ / 10% glycerol / 0.2 mM PMSF) at the concentration of 1 × 10^6^ cells/ml. The cell suspension was incubated 20 min on ice and mixed by vortexing. After 5 min centrifugation at 4000 rpm at 4°C, the pellet was washed twice in PBS and resuspended in Buffer C (20 mM Hepes pH 7.9 / 300 mM NaCl / 0.2% NP-40 / 25% glycerol / 1 mM MgCl_2_/ 0.2 mM PMSF / 1 x Protease inhibitor mix / 0.3 mM DTT) at the concentration of 1 x 10^6^ nuclei/100 μl. After 20 min of incubation on ice, the nuclei suspension was centrifuged 5 min at 13200 rpm at 4°C and the supernatant was collected.

The NE from MT cells overexpressing the RBPJ constructs was prepared as follows. Briefly, 10 × 10^6^ cells were washed with PBS and resuspended in 200 μl of Buffer 1 (10 mM Hepes pH 7.9 / 10 mM KCl / 0.1 mM EDTA / 0.1 mM EGTA / 1 mM ßME, supplemented with PMSF). The cell suspension was incubated 10 min on ice, 5 μl of 10% NP-40 were added and mixed by vortexing. After 10 sec of centrifugation at 13000 rpm at 4°C, the nuclei pellet was washed twice in 500 μl of Buffer 1 and resuspended in 100 μl of Buffer 2 (20 mM Hepes pH 7.9 / 400 mM NaCl / 1 mM EDTA / 1 mM EGTA / 1 mM ßME, supplemented with PMSF). After 20 min of incubation on ice, the nuclei suspension was centrifuged 10 min at 13000 rpm at 4°C and the supernatant was collected for further analysis.

Protein concentration was measured by Bradford assay (Sigma-Aldrich) and samples were boiled after adding SDS-polyacrylamide gel loading buffer. Samples were resolved by SDS-Page and analyzed by Western blotting using antibodies against GAPDH (abcam, ab8245), GFP (Roche, 11814460001) or TBP (Santa Cruz, sc-273). Briefly, membranes were blocked in 5% milk, 1x TBS, 0.1% Tween-20 (TBS-T) and primary antibodies were diluted in 5% milk, TBS-T. After incubation over night at 4°C, membranes were washed in TBS-T, secondary antibodies against mouse (Cell Signaling, #7076S) or rabbit (Cell Signaling, #7074S) were diluted in 5% milk TBS-T and finally membranes were washed in TBS-T.

In the case of the RBPJ Western blotting the procedure was as follows. Briefly, membranes were blocked in 5% milk, 1x TBS and the RBPJ antibody (Cosmo Bio Co. LTD, 2ZRBP2) was diluted 1:1000 in 5% BSA, 1x TBS, 0.3% NP40. After incubation over night at 4°C, membranes were washed three times 15 min each in 1x TBS / 0.5 M NaCl / 0.5% Triton X-100 and the secondary antibody against rat (Jackson ImmunoResearch, 112–035–072) was diluted 1:5000 in 5% BSA, 1x TBS, 0.3% NP-40. Membranes were washed three times 15 min each in 1x TBS / 0.5 M NaCl / 0.5% Triton X-100. All membranes were finally incubated with ECL solution and chemiluminescence was detected with a light sensitive film.

## Acknowledgements

We thank Roswitha Rittelmann, Sabine Schirmer, and Thomas Schmidt-Wöll for excellent technical assistance. This work was supported by National Institutes of Health Grants 5R01CA178974 (RAK). It was also supported by the collaborative research grant TRR81 and the grant (BO 1639/9–1) by the DFG (German Research Foundation) and the Excellence Cluster for Cardio Pulmonary System (ECCPS) in Giessen to TB and the University Medical Center Giessen and Marburg (UKGM to BG). This work was further supported by German Research Foundation (DFG) Grant SFB1074/A3 and by the BMBF (Federal Ministry of Education and Research, research nucleus SyStAR) to FO. LP is supported by the DFG (GRK 2253 - HEIST). This research used resources of the Advanced Photon Source, a U.S. Department of Energy (DOE) Office of Science User Facility operated for the DOE Office of Science by Argonne National Laboratory under Contract No. DE-AC02–06CH11357.

---

**Table S1.**
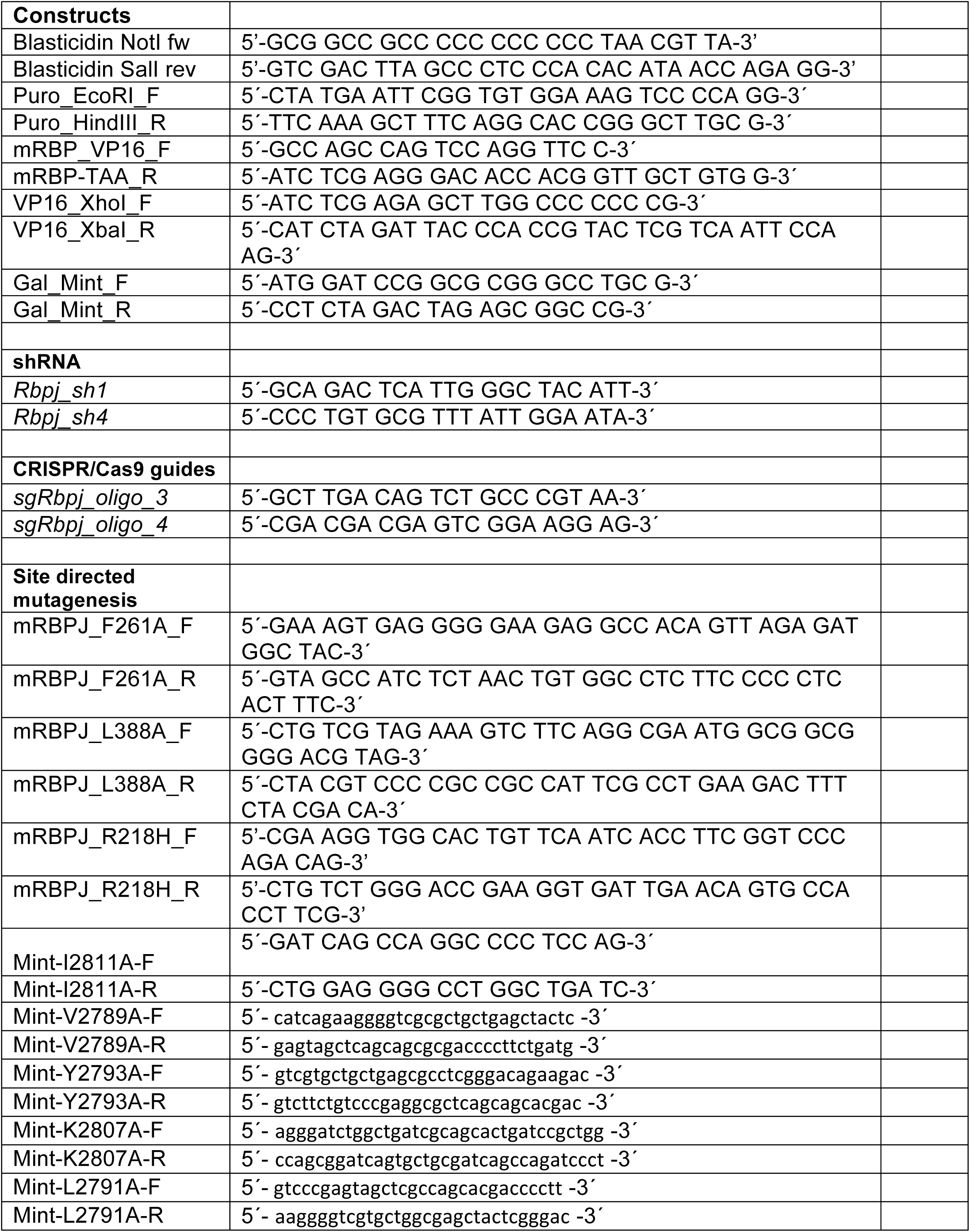
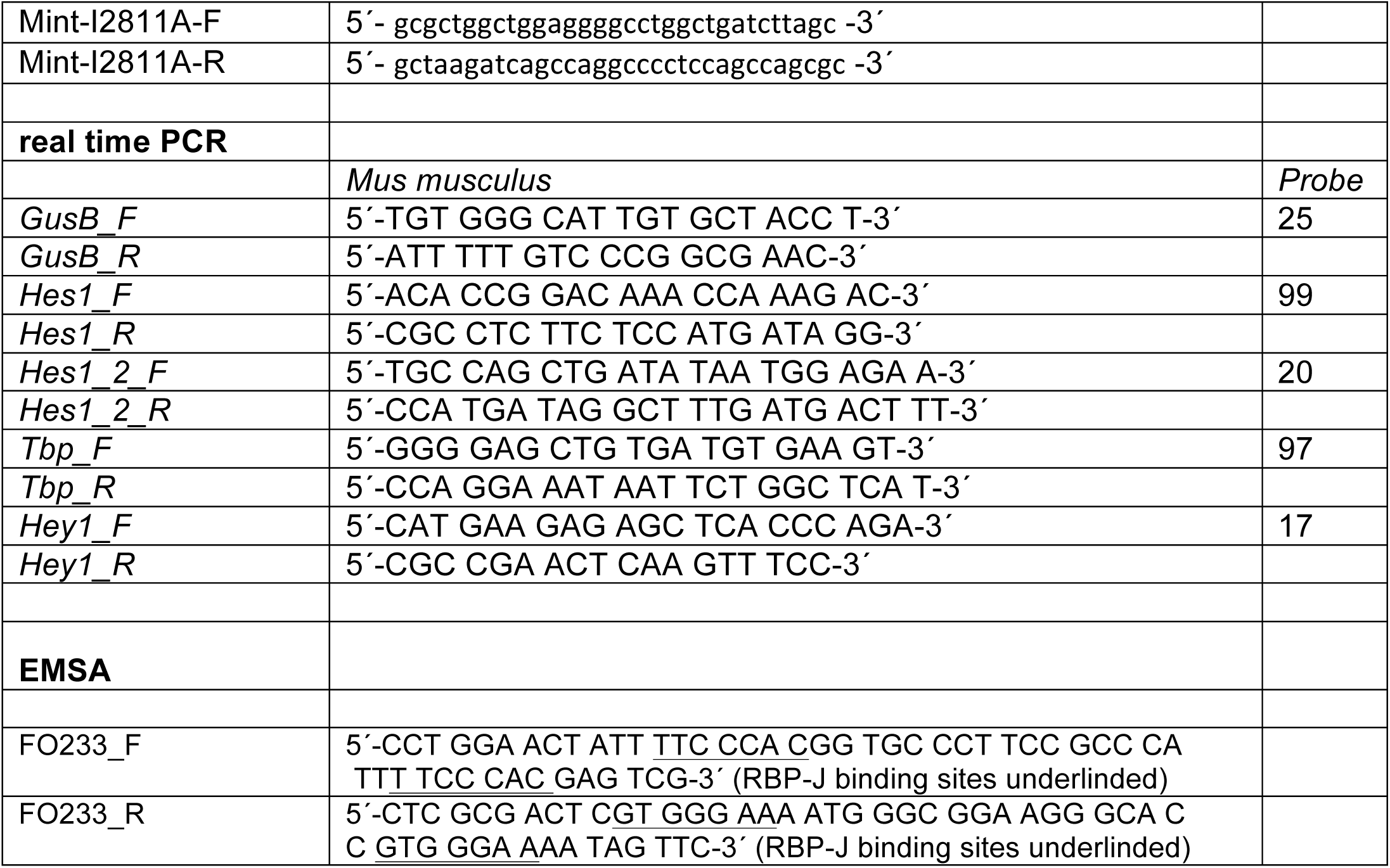
Oligonucleotides used for plasmid construction, EMSAs, shRNAs, CRISPR/Cas9, and qPCR experiments.

**Table S2.** Constructs cloned by PCR.

| Plasmid | Template | Primer Forward | Primer Reverse | Enzymes | Ligation |
| --- | --- | --- | --- | --- | --- |
| pMIGR1<br>pSV40-Puro | pLXSPHAMEl18<br>ATG | Puro EcoRI F | Puro HindIII R | EcoRI/ HindIII | pMIGR1 IRES-GFP |
| pcDNA3.1.Flag<br>-2-mRBPJ | pcDNA3.1.Flag-2-mRBPJ | mRBP_VP16_<br>F | mRBP-TAA_R | PshAI/XhoI | pcDNA3.1.Flag<br>-2-mRBPJΔstop |
| pcDNA3.1.Flag<br>-2-mRBPJΔstop | pFA-CMV | VP16_XhoI_F | VP16_XbaI_R | XhoI/XbaI | pcDNA3.1-Flag-2-mRBPJ-VP16 |
| pFA-CMV | pcDNA3.1_HisC<br>_MINT_2776-2833 | Gal-Mint_F | Gal-Mint_R | BamHI/XbaI | pFA-CMV-Mint-2776-2833(wt) |
| pMY-Bio-IRES-Blasticidin | pMSCV-FLAG-hRBP-J Ires<br>Blasticidin | Blasticidin NotI<br>fw | Blasticidin Sall<br>rev | NotI/Sall | pMY BioTip60<br>IRES-GFP |

**Figure S1.** <u>Composition and analysis of the asymmetric unit of the RBPJ/MBP-SHARP/DNA complex crystals</u>. (A) The asymmetric unit of the RBPJ/MBP-SHARP/DNA complex crystals contains two RBPJ/MBP-SHARP/DNA complexes. RBPJ, MBP, and SHARP are shown as ribbon diagrams and colored green, cyan, and purple, respectively. The DNA is colored gray. (B) ITC thermogram showing the affinity of MBP-SHARP for RBPJ. (C-D) Structural overlays of the two complexes contained within the asymmetric unit of the crystals. (C) Structural alignment overall all cα atoms. (D) Resulting structural overlay when only RBPJ cα atoms are aligned. (E) Resulting structural overlay when only MBP cα atoms are aligned. (F) Resulting structural overlay when only SHARP cα atoms are aligned. Red arrows highlight the large conformational differences in the poorly ordered linkers of SHARP that lie between the regions that bind the BTD and CTD of RBPJ.

**Figure S2.** <u>Control studies of the structure based RBPJ mutants.</u>. (A) Differential scanning fluorimetry (DSF) was used to analyze the melting temperatures (Tm) of wild-type and mutant RBPJ proteins. Figure shows melt curve and derivative plot. Tm determined for wild-type and mutant RBPJ proteins is the mean ± SD of experiments performed in triplicate. (B) Upper panel, analysis of DNA binding by EMSA of wild-type RBPJ and select RBPJ mutants. Oligomeric duplex DNA probe used in EMSA 5'-CCTGGAACTATT**<u>TTCCCACG</u>**GTGCCCTTCCGCCCATT**<u>TTCCCACG</u>**AGTCG-3' with RBPJ binding sites in bold and underlined. The DNA interaction bands are labeled A and B. The asterisk denotes a nonspecific background band. Addition of α-Flag antibodies lead to supershifted Flag-RBPJ/DNA complexes (complexes a and b) in the EMSA, demonstrating specificity. Lower panel, N-terminally tagged Flag-RBPJ fusion proteins were translated *in vitro* with the TNT T7 coupled Reticulocyte Lysate System Kit (Promega) and relative protein levels were analyzed by Western blotting. Flag-RBPJ/DNA complexes, labeled a and b, were supershifted in the EMSA using α-Flag antibodies, demonstrating specificity. (C) Subcellular localization of RBPJ proteins after expression in HeLa cells. HeLa cells were transiently transfected with 0.3 μg of the respective plasmids GFP-RBPJ: WT, F261A, L388A, and F261A/L388A. After 24 h cells were fixed and the images were taken 24 h after transfection with DAPI staining under the fluorescence microscope using a 63x lens. The superimposition (right column) of the DAPI signal (center) with the GFP signal (left) illustrates the predominantly nuclear localization of the RBPJ proteins. (D) NICD binding of wild-type and mutant RBPJ proteins from cells. HEK293 cells were transfected with the plasmids GFP-RBPJ^WT^, GFP-RBPJ^F261A^ GFP-RBPJ^L388A^, or GFP-RBPJ^F261A/L388A^ plasmids together with with Flag-NICD (lanes 2–6). Immunoprecipitation was performed with the α-Flag antibody agarose beads and detected by Western blotting using an α-GFP antibody. The expression of the Flag-tagged NICD protein (lower blot) could be detected using an α-Flag antibody. The asterisk in the upper blot marks the heavy chain of the antibody used for immunoprecipitation.

## References

1 Kovall, R. A., Gebelein, B., Sprinzak, D. & Kopan, R. The Canonical Notch Signaling Pathway: Structural and Biochemical Insights into Shape, Sugar, and Force. Dev Cell 41, 228–241, doi:10.1016/j.devcel.2017.04.001 (2017).

2 Bray, S. J. Notch signalling in context. Nat Rev Mol Cell Biol 17, 722–735, doi:10.1038/nrm.2016.94 (2016).

3 Siebel, C. & Lendahl, U. Notch Signaling in Development, Tissue Homeostasis, and Disease. Physiological reviews 97, 1235–1294, doi:10.1152/physrev.00005.2017 (2017).

4 Braune, E. B. & Lendahl, U. Notch -- a goldilocks signaling pathway in disease and cancer therapy. Discovery medicine 21, 189–196 (2016).

5 Ntziachristos, P., Lim, J. S., Sage, J. & Aifantis, I. From fly wings to targeted cancer therapies: a centennial for notch signaling. Cancer cell 25, 318–334, doi:10.1016/j.ccr.2014.02.018 (2014).

6 Kovall, R. A., & Blacklow, S. C. Mechanistic insights into Notch receptor signaling from structural and biochemical studies. Curr Top Dev Biol 92, 31–71, doi:S0070-2153(10)92002-4 [pii] 10.1016/S0070-2153(10)92002-4 (2010).

7 Borggrefe, T. & Oswald, F. The Notch signaling pathway: transcriptional regulation at Notch target genes. Cell Mol Life Sci 66, 1631–1646 (2009).

8 Kuroda, K. et al. Regulation of marginal zone B cell development by MINT, a suppressor of Notch/RBP-J signaling pathway. Immunity 18, 301–312 (2003).

9 Oswald, F. et al. SHARP is a novel component of the Notch/RBP-Jkappa signalling pathway. Embo J 21, 5417–5426 (2002).

10 Maier, D. Hairless: the ignored antagonist of the Notch signalling pathway. Hereditas 143, 212–221, doi:10.1111/j.2007.0018-0661.01971.x (2006).

11 Taniguchi, Y., Furukawa, T., Tun, T., Han, H. & Honjo, T. LIM protein KyoT2 negatively regulates transcription by association with the RBP-J DNA-binding protein. Mol Cell Biol 18, 644–654 (1998).

12 Xu, T. et al. RBPJ/CBF1 interacts with L3MBTL3/MBT1 to promote repression of Notch signaling via histone demethylase KDM1A/LSD1. EMBO J 36, 3232–3249, doi:10.15252/embj.201796525 (2017).

13 Tabaja, N., Yuan, Z., Oswald, F. & Kovall, R. A. Structure-function analysis of RBP-J-interacting and tubulin-associated (RITA) reveals regions critical for repression of Notch target genes. J Biol Chem 292, 10549–10563, doi:10.1074/jbc.M117.791707 (2017).

14 Wacker, S. A. et al. RITA, a novel modulator of Notch signalling, acts via nuclear export of RBP-J. Embo J 30, 43–56, doi:emboj2010289 [pii] 10.1038/emboj.2010.289 (2011).

15 Wilson, J. J. & Kovall, R. A. Crystal structure of the CSL-Notch-Mastermind ternary complex bound to DNA. Cell 124, 985–996, doi:10.1016/j.cell.2006.01.035 (2006).

16 Nam, Y., Sliz, P., Song, L., Aster, J. C. & Blacklow, S. C. Structural basis for cooperativity in recruitment of MAML coactivators to Notch transcription complexes. Cell 124, 973–983, doi:10.1016/j.cell.2005.12.037 (2006).

17 Kovall, R. A. & Hendrickson, W. A. Crystal structure of the nuclear effector of Notch signaling, CSL, bound to DNA. EMBO J 23, 3441–3451, doi:10.1038/sj.emboj.7600349 (2004).

18 Yuan, Z. et al. Structure and Function of the Su(H)-Hairless Repressor Complex, the Major Antagonist of Notch Signaling in Drosophila melanogaster. PLoS Biol 14, e1002509, doi:10.1371/journal.pbio.1002509 (2016).

19 Collins, K. J., Yuan, Z. & Kovall, R. A. Structure and function of the CSL-KyoT2 corepressor complex: a negative regulator of Notch signaling. Structure 22, 70–81, doi:10.1016/j.str.2013.10.010 (2014).

20 Shi, Y. et al. Sharp, an inducible cofactor that integrates nuclear receptor repression and activation. Genes Dev 15, 1140–1151, doi:10.1101/gad.871201 (2001).

21 Newberry, E. P., Latifi, T. & Towler, D. A. The RRM domain of MINT, a novel Msx2 binding protein, recognizes and regulates the rat osteocalcin promoter. Biochemistry 38, 10678–10690, doi:10.1021/bi990967j (1999).

22 Salat, D., Liefke, R., Wiedenmann, J., Borggrefe, T. & Oswald, F. ETO, but not leukemogenic fusion protein AML1/ETO, augments RBP-Jkappa/SHARP-mediated repression of notch target genes. Mol Cell Biol 28, 3502–3512, doi:10.1128/MCB.01966-07 (2008).

23 Oswald, F. et al. RBP-Jkappa/SHARP recruits CtIP/CtBP corepressors to silence Notch target genes. Mol Cell Biol 25, 10379–10390, doi:10.1128/MCB.25.23.10379-10390.2005 (2005).

24 Oswald, F. et al. A phospho-dependent mechanism involving NCoR and KMT2D controls a permissive chromatin state at Notch target genes. Nucleic Acids Res 44, 4703–4720 (2016).

25 McHugh, C. A. et al. The Xist lncRNA interacts directly with SHARP to silence transcription through HDAC3. Nature 521, 232–236, doi:10.1038/nature14443 (2015).

26 Chu, C. et al. Systematic discovery of Xist RNA binding proteins. Cell 161, 404–416, doi:10.1016/j.cell.2015.03.025 (2015).

27 VanderWielen, B. D., Yuan, Z., Friedmann, D. R. & Kovall, R. A. Transcriptional repression in the Notch pathway: thermodynamic characterization of CSL-MINT (Msx2-interacting nuclear target protein) complexes. J Biol Chem 286, 14892–14902, doi:10.1074/jbc.M110.181156 (2011).

28 Moon, A. F., Mueller, G. A., Zhong, X. & Pedersen, L. C. A synergistic approach to protein crystallization: combination of a fixed-arm carrier with surface entropy reduction. Protein Sci 19, 901–913, doi:10.1002/pro.368 (2010).

29 Choi, S. H. et al. Conformational locking upon cooperative assembly of notch transcription complexes. Structure 20, 340–349, doi:10.1016/j.str.2011.12.011 (2012).

30 Friedmann, D. R., Wilson, J. J. & Kovall, R. A. RAM-induced allostery facilitates assembly of a notch pathway active transcription complex. J Biol Chem 283, 14781–14791, doi:M709501200 [pii] 10.1074/jbc.M709501200 (2008).

31 Johnson, S. E., Ilagan, M. X., Kopan, R. & Barrick, D. Thermodynamic analysis of the CSL x Notch interaction: distribution of binding energy of the Notch RAM region to the CSL beta-trefoil domain and the mode of competition with the viral transactivator EBNA2. J Biol Chem 285, 6681–6692, doi:10.1074/jbc.M109.019968 (2010).

32 Del Bianco, C., Aster, J. C. & Blacklow, S. C. Mutational and energetic studies of Notch 1 transcription complexes. J Mol Biol 376, 131–140, doi:S0022-2836(07)01560-4 [pii] 10.1016/j.jmb.2007.11.061 (2008).

33 Lubman, O. Y., Ilagan, M. X., Kopan, R. & Barrick, D. Quantitative dissection of the Notch:CSL interaction: insights into the Notch-mediated transcriptional switch. J Mol Biol 365, 577–589, doi:10.1016/j.jmb.2006.09.071 (2007).

34 Maier, D. et al. Structural and functional analysis of the repressor complex in the Notch signaling pathway of Drosophila melanogaster. Mol Biol Cell 22, 3242–3252, doi:mbc.E11-05-0420 [pii] 10.1091/mbc.E11-05-0420 (2011).

35 Kurth, P., Preiss, A., Kovall, R. A. & Maier, D. Molecular analysis of the notch repressor-complex in Drosophila: characterization of potential hairless binding sites on suppressor of hairless. PLoS One 6, e27986, doi:10.1371/journal.pone.0027986 PONE-D-11-18755 [pii] (2011).

36 Yuan, Z., Friedmann, D. R., VanderWielen, B. D., Collins, K. J. & Kovall, R. A. Characterization of CSL (CBF-1, Su(H), Lag-1) mutants reveals differences in signaling mediated by Notch1 and Notch2. J Biol Chem 287, 34904–34916, doi:M112.403287 [pii] 10.1074/jbc.M112.403287 (2012).

37 Jung, C., Mittler, G., Oswald, F. & Borggrefe, T. RNA helicase Ddx5 and the noncoding RNA SRA act as coactivators in the Notch signaling pathway. Biochimica et biophysica acta 1833, 1180–1189, doi:10.1016/j.bbamcr.2013.01.032 (2013).

38 Bray, S. & Bernard, F. Notch targets and their regulation. Curr Top Dev Biol 92, 253–275, doi:S0070-2153(10)92008-5 [pii] 10.1016/S0070-2153(10)92008-5 (2010).

39 Surendran, K. et al. The contribution of Notch1 to nephron segmentation in the developing kidney is revealed in a sensitized Notch2 background and can be augmented by reducing Mint dosage. Developmental biology 337, 386–395, doi:10.1016/j.ydbio.2009.11.017 (2010).

40 Tsuji, M., Shinkura, R., Kuroda, K., Yabe, D. & Honjo, T. Msx2-interacting nuclear target protein (Mint) deficiency reveals negative regulation of early thymocyte differentiation by Notch/RBP-J signaling. Proc Natl Acad Sci USA 104, 1610–1615, doi:10.1073/pnas.0610520104 (2007).

41 Chan, S. K. K. et al. Role of co-repressor genomic landscapes in shaping the Notch response. PLoS genetics 13, e1007096, doi:10.1371/journal.pgen.1007096 (2017).

42 Castel, D. et al. Dynamic binding of RBPJ is determined by Notch signaling status. Genes Dev 27, 1059–1071, doi:10.1101/gad.211912.112 (2013).

43 Kao, H. Y. et al. A histone deacetylase corepressor complex regulates the Notch signal transduction pathway. Genes Dev 12, 2269–2277 (1998).

44 Hsieh, J. J. & Hayward, S. D. Masking of the CBF1/RBPJ kappa transcriptional repression domain by Epstein-Barr virus EBNA2. Science 268, 560–563 (1995).

45 Torella, R. et al. A combination of computational and experimental approaches identifies DNA sequence constraints associated with target site binding specificity of the transcription factor CSL. Nucleic Acids Res 42, 10550–10563, doi:10.1093/nar/gku730 (2014).

46 Friedmann, D. R., & Kovall, R. A. Thermodynamic and structural insights into CSL-DNA complexes. Protein Sci 19, 34–46, doi:10.1002/pro.280 (2010).

47 Gomez-Lamarca, M. J. et al. Activation of the Notch Signaling Pathway In Vivo Elicits Changes in CSL Nuclear Dynamics. Dev Cell 44, 611–623 e617, doi:10.1016/j.devcel.2018.01.020 (2018).

48 Ariyoshi, M. & Schwabe, J. W. A conserved structural motif reveals the essential transcriptional repression function of Spen proteins and their role in developmental signaling. Genes Dev 17, 1909–1920 (2003).

49 Zehender, A. et al. Conservation of the Notch antagonist Hairless in arthropods: functional analysis of the crustacean Daphnia pulex Hairless gene. Development genes and evolution 227, 339–353, doi:10.1007/s00427-017-0593-4 (2017).

50 McCoy, A. J. et al. Phaser crystallographic software. Journal of applied crystallography 40, 658–674, doi:10.1107/S0021889807021206 (2007).

51 Adams, P. D. et al. PHENIX: a comprehensive Python-based system for macromolecular structure solution. Acta crystallographica. Section D, Biological crystallography 66, 213–221, doi:10.1107/S0907444909052925 (2010).

52 Emsley, P. & Cowtan, K. Coot: model-building tools for molecular graphics. Acta crystallographica. Section D, Biological crystallography 60, 2126–2132 (2004).

53 Smart, O. S. et al. Exploiting structure similarity in refinement: automated NCS and target-structure restraints in BUSTER. Acta crystallographica. Section D, Biological crystallography 68, 368–380, doi:10.1107/S0907444911056058 (2012).

54 Davis, I. W. et al. MolProbity: all-atom contacts and structure validation for proteins and nucleic acids. Nucleic Acids Res 35, W375–383 (2007).

55 Krissinel, E. & Henrick, K. Inference of macromolecular assemblies from crystalline state. J Mol Biol 372, 774–797 (2007).

